# ULK complex organization in autophagy by a C-shaped FIP200 N-terminal domain dimer

**DOI:** 10.1101/840009

**Authors:** Xiaoshan Shi, Adam L. Yokom, Chunxin Wang, Lindsey N. Young, Richard J. Youle, James H. Hurley

## Abstract

The autophagy-initiating human ULK complex consists of the kinase ULK1/2, FIP200, ATG13, and ATG101. Hydrogen-deuterium exchange mass spectrometry was used to map their mutual interactions. The N-terminal 640 residues (NTD) of FIP200 interact with the C-terminal IDR of ATG13. Mutations in these regions abolish their interaction. Negative stain electron microscopy (EM) and multiangle light scattering showed that FIP200 is a dimer whilst a single molecule each of the other subunits is present. The FIP200 NTD is flexible in the absence of ATG13, but in its presence adopts the shape of the letter C ~20 nm across. The ULK1 EAT domain interacts loosely with the NTD dimer, while the ATG13-ATG101 HORMA dimer does not contact the NTD. Cryo-EM of the NTD dimer revealed a structure similarity to the scaffold domain of TBK1, suggesting an evolutionary similarity between the autophagy initiating TBK1 kinase and the ULK1 kinase complex.

**Summary:** The human ULK complex consists of ULK1/2, FIP200, ATG13, and ATG101. We found that the FIP200 N-terminal domain is a C-shaped dimer that binds directly to a single ATG13 molecule and serves as the organizing hub of the complex.

## Introduction

Macroautophagy (henceforward, autophagy) is the conserved eukaryotic cellular process responsible for replenishment of biosynthetic precursors during starvation (Wen and Klionsky, 2016), and engulfment and degradation of molecular aggregates, organelles, intracellular pathogens, and many other cellular substrates (Anding and Baehrecke, 2017; Gomes and Dikic, 2014; Zaffagnini and Martens, 2016). Autophagy proceeds by the *de novo* formation of a cup-shaped double membrane known as the phagophore or isolation membrane. The phagophore double membrane grows such that it engulfs and isolates its substrates. Upon sealing of the double membrane, the mature structure is referred to as an autophagosome. The autophagosome then fuses with the lysosome, leading to the degradation of the material within the autophagosome. The proteins and protein complexes responsible for these steps have been identified (Mizushima et al., 2011). In mammals (Bento et al., 2016), these include the ULK1 (unc-51 like autophagy activating kinase 1) protein kinase complex, the class III phosphatidylinositol 3-kinase complexes (PI3KC3), the phosphatidylinositol 3-phosphate (PI(3)P)-sensing WIPI proteins, the lipid transporter ATG2 (autophagy related 2), the integral membrane protein ATG9, the ubiquitin-like ATG8 proteins, machinery for conjugating ATG8 protein to lipid membranes, the autophagy adaptors that connect substrates to the ULK1 complex and to ATG8 proteins, and TBK1 (TANK binding kinase 1), which phosphoregulates autophagy adaptors. The mechanisms by which these protein complexes orchestrate autophagosome initiation, growth, closure, and delivery to the lysosome are being actively sought (Hurley and Young, 2017; Mercer et al., 2018).

The mammalian ULK1 complex is the most upstream of the core protein complexes that make autophagosomes (Itakura and Mizushima, 2010; Karanasios et al., 2013; Karanasios et al., 2016; Koyama-Honda et al., 2013). It is the mammalian counterpart of the yeast Atg1 complex, whose assembly is the main trigger for starvation-induced autophagy in yeast (Kamada et al., 2000). In starvation and TORC1 inhibition, the yeast Atg1 complex assembles from protein kinase Atg1, the bridging subunit Atg13, and the constitutively assembled scaffold Atg17-Atg29-Atg31 to initiate the phagophore (Kamada et al., 2010). The structure of Atg17 has the form of an S-shaped dimer (Ragusa et al., 2012) whose dimensions and curvature are suited to promoting cup-shaped membrane structures (Bahrami et al., 2017). In yeast, there are starvation and TORC1-independent forms of selective autophagy that use Atg1 and Atg13, but with the Atg11 scaffold replacing the Atg17 subcomplex (Yorimitsu and Klionsky, 2005).

The ULK1 complex consists of ULK1 itself, the scaffolding subunit FIP200 (FAK family kinase-interacting protein of 200 kDa; also known as RB1-inducible coiled-coil protein 1 (RB1CC1)), ATG13 and ATG101 (Ganley et al., 2009; Hosokawa et al., 2009; Jung et al., 2009; Mercer et al., 2009). ULK1 can in most cases be replaced by its paralog ULK2, a closely related serine/threonine kinase which is partially interchangeable within the ULK complex (Mizushima, 2010). ULK1 and ULK2 are the mammalian paralogs of Atg1. ULK1 contains the only known catalytic activity within the complex. The crystal structure of its N-terminal kinase domain is known (Lazarus et al., 2015). The ULK1 kinase targets downstream autophagic machinery including ATG14, VPS34, ATG9, ATG4, and others (Papinski and Kraft, 2016; Zachari and Ganley, 2017). ULK1 contains a C-terminal early autophagy targeting/tethering (EAT) domain, which is connected to the kinase domain by a ~550 residue long intrinsically disordered region (IDR) and targets ULK1 by binding to a motif in the C-terminus of ATG13 (Chan et al., 2009; Hieke et al., 2015).

ATG13 consists of an N-terminal HORMA (Hop/Rev7/Mad2) domain, which dimerizes with the HORMA domain of ATG101 (Qi et al., 2015; Suzuki et al., 2015) and a long C-terminal IDR which binds to FIP200 (Jung et al., 2009; Wallot-Hieke et al., 2018) and ULK1 (Hieke et al., 2015; Wallot-Hieke et al., 2018). The ATG101 HORMA contains an exposed WF finger motif that is important for autophagy(Suzuki et al., 2015). It is not clear what the interaction partner of the WF finger is or how the HORMA dimer fits into the larger organization of the complex.

FIP200 is comprised of 1594 residues, is essential for autophagy, and is considered the functional counterpart of the yeast Atg11 and Atg17 scaffold subunits (Hara et al., 2008) (Fig. 1A). FIP200, however, has no sequence homology to Atg11 and Atg17 apart from the C-terminal 100-residue CLAW domain of FIP200 and Atg11. This is the only portion of FIP200 whose structure is known (Turco et al., 2019). The remainder of FIP200 consists of a ~640 residue N-terminal domain, followed by an IDR linker and a coiled-coil domain comprising ~750 residues. Targeting of FIP200 by the autophagy adaptors NDP52 or p62 to mitochondria (Vargas et al., 2019), *Salmonella* (Ravenhill et al., 2019), or ubiquitinated cargo (Turco et al., 2019) condensates triggers phagophore initiation leading to their engulfment. Thus, FIP200 is absolutely central to autophagy initiation. Yet as one of the largest proteins in the autophagic machinery, FIP200 has been among the most difficult to study *in vitro*. The lack of reported motifs or sequence homology in the N-terminal 1500 residues has also slowed progress in understanding this critical part of the autophagy machinery.

**Figure 1.**
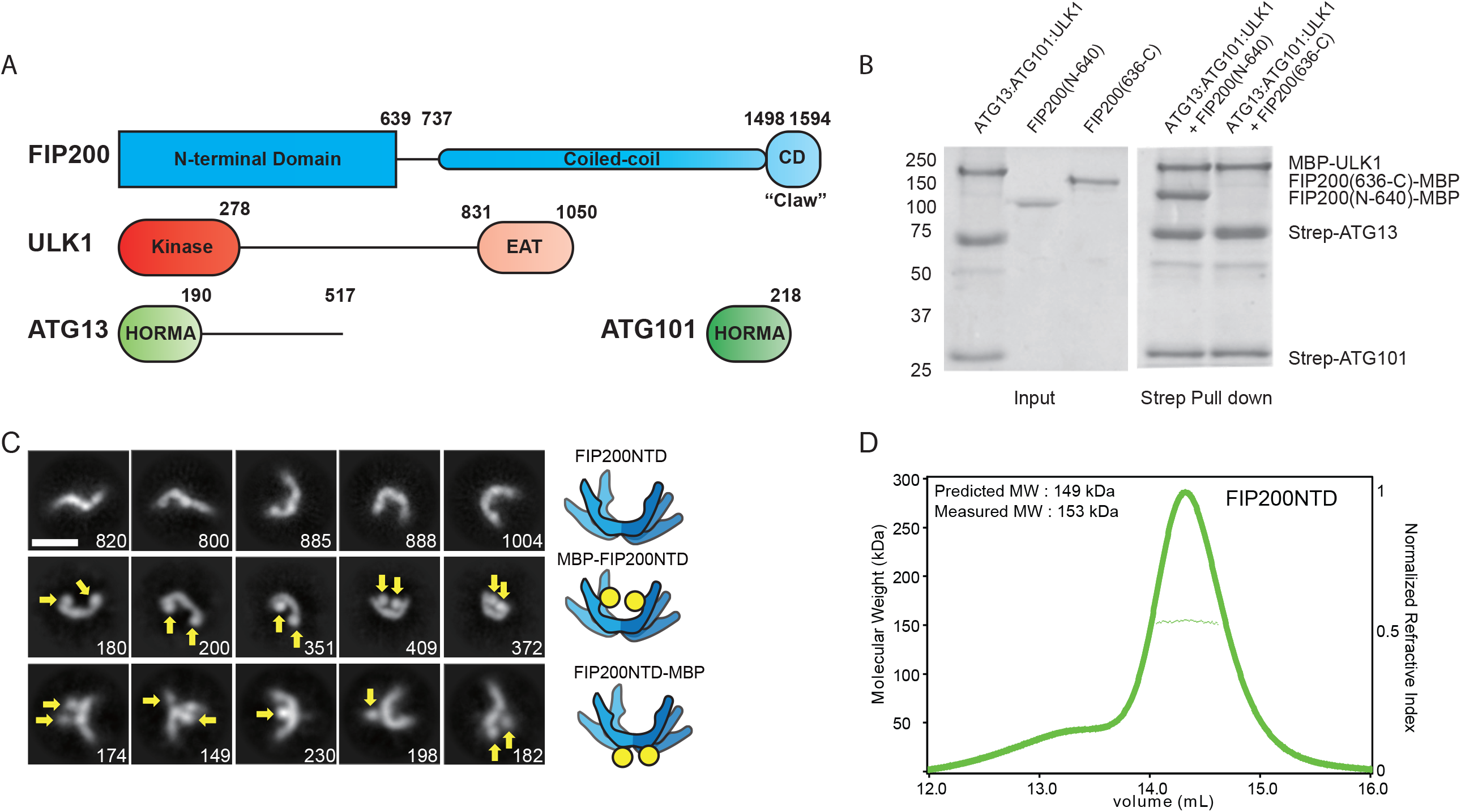
The FIP200 NTD scaffolds the ULK Complex as a homodimer. (A) Domain diagram of the ULK complex proteins. (B) Pull-down assay of ULK1, ATG13 and ATG101 with FIP200 NTD (N-640) and FIP200 CTD (636-C). Strep-Tactin resin was loaded with MBP-ULK1:Strep-ATG13:Strep-ATG101 complex to pull down FIP200(N-640)-MBP and FIP200(636-C)-MBP. The pull-down results were visualized by SDS-PAGE and Coomassie blue staining. (C) Negative Stain EM 2D class averages for FIP200 NTD (top), N terminal MBP tagged FIP200 NTD (middle) and C-terminal MBP tagged FIP200 NTD (bottom). Densities corresponding to MBP tags are labeled with yellow arrows. Scale bar is 20 nm. (D) Multi-angle light scattering and size exclusion chromatography trace of FIP200 NTD shows the predicted and measured MW of the dimeric FIP200 NTD.

In this study, we expressed and purified the human ULK1 complex in order to understand its structural organization. It became clear early in these studies that the full ULK1 complex, with its extensive IDR content and 750-residue FIP200 coiled coil, and dissociable interactions between most of the subunits, is not a typical well-ordered, co-assembled constitutive complex. Its dynamic character makes it exceptionally challenging for structural studies. Nevertheless, we were able to use hydrogen-deuterium exchange coupled to mass spectrometry (HDX-MS), electron microscopy (EM) and MALS to map the organization of the complex. These data show that the N-terminal domain of FIP200 serves a C-shaped dimeric hub for assembly of ULK1 complex.

## Results

### FIP200 NTD assembles with ULK1, ATG13, and ATG101

Given that FIP200 is essential for autophagy and is the largest protein in ULK1 complex, we began with the hypothesis that some part of FIP200 was likely to be the main hub organizing the complex. We sought to identify the minimal domain needed for the assembly of the core ULK1 complex. Both the N-terminal (NTD, 1-640) and C-terminal (CTD, 636-1594) domains (Fig. 1A) of FIP200 were expressed in HEK 293T cells as N-terminal GST and C-terminal MBP fusions. The remaining three subunits, ULK1, ATG13, and ATG101 were separately co-expressed with each other and purified. Purified FIP200 NTD-MBP and FIP200 CTD-MBP were used in pulldown assay with the ternary ULK1:ATG13:ATG101 complex. These experiments showed there to be an interaction between ULK1:ATG13:ATG101 and the FIP200 NTD, but not the CTD (Fig. 1B).

### The FIP200 NTD is a C-shaped dimer

We characterized the overall size and shape of FIP200 NTD using negative stain electron microscopy (NSEM) and multiangle light scattering (MALS). NSEM 2D classification of FIP200 NTD showed a variety of shapes (Fig. 1C, Fig. S1A-C, Table S1), with maximum dimensions ranging from 10 to 24 nm. Many of the 2D averages were in the shape of the letter C. Others resembled singly bent rods or S-shapes (Fig. 1C). MBP tags were fused to either the N- or C-terminus in order to mark the location of each end by the presence of additional density compared to untagged FIP200 NTD. 2D class averages of the MBP N-terminal tagged construct displayed a similar variety of shapes, with two additional densities corresponding to two MBP tags at the tips of the density (Fig. 1C, see arrows). This observation suggested that FIP200 NTD is a dimer, and that the N-termini are distal to the dimer interface. The C-terminal MBP tags were localized near the center of the C-shape. The C-terminal tags displayed more dispersed positions, with either one or two tags visualized. Together, the MBP tags suggest the FIP200 NTD forms a dimeric structure with the C-termini close to the dimer interface and the N-termini at the tips.

To assess the oligomeric state of FIP200 by an independent technique, we used size exclusion chromatography (SEC) coupled to multiangle light scattering (MALS) to analyze the molecular weight of FIP200 NTD in solution. The SEC-MALS resulted in a single peak with a molecular weight of 153 kDa (Fig. 1D). This value corresponds closely to the predicted MW of 149kDa of the FIP200 NTD dimer.

### Mapping FIP200 NTD interactions with the rest of ULK1 the complex

We sought to determine the minimal region(s) of ULK1:ATG13:ATG101 interacting with FIP200 NTD. Strep-ATG13:Strep-ATG101 was efficiently pulled down by FIP200NTD alone, but MBP-ULK1 was not (Fig. 2A). Pull down of ULK1 was recovered in the presence of ATG13:ATG101 (Fig. 2A). This shows that ATG13:ATG101 interacts directly FIP200 NTD, whilst ULK1 recruitment to FIP200 NTD depends on the presence of ATG13:ATG101. We characterized the minimal FIP200 NTD:ATG13:ATG101 subcomplex by NSEM. 2D classification showed distinct C shapes, as seen for FIP200 NTD alone (Fig. 2B, Fig. S1D-F, Table S1). However, we did not observe any of the bent rods and S-shapes seen in the absence of ATG13:ATG101. Thus ATG13:ATG101 stabilizes the C-shaped conformation of FIP200 NTD. No extra density corresponding to HORMA dimers was seen in the 2D averages, suggesting that the position of the HORMA dimer is not ordered with respect the FIP200 NTD.

**Figure 2.**
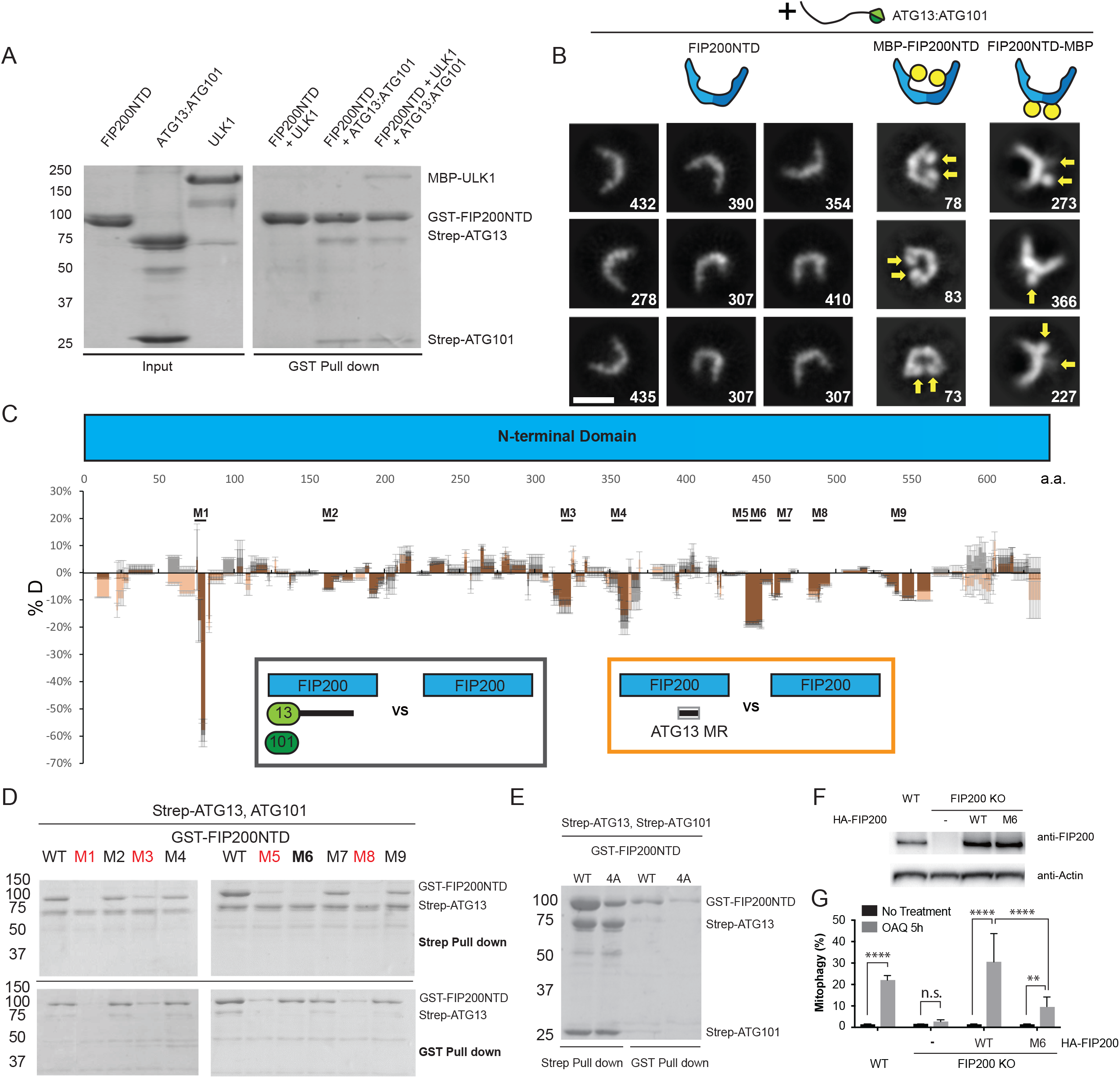
Mapping the FIP200 NTD interaction sites with ATG13 and ATG101. (A) Pull-down assay of FIP200 NTD with ULK1, ATG13:ATG101 and both. GSH resin was loaded with GST-FIP200NTD to pull down MBP-ULK1, Strep-ATG13:Strep-ATG101 and both. The pull-down results were visualized by SDS-PAGE and Coomassie blue staining. (B) Negative stain EM 2D class averages of FIP200 NTD:ATG13:ATG101, MBP-FIP200 NTD:ATG13:ATG101 and FIP200 NTD-MBP:ATG13:ATG101. Densities corresponding to MBP tags are labeled with yellow arrows. Scale bar is 20 nm. (C) Difference of Hydrogen Deuterium Exchange percentages of the FIP200 NTD alone vs FIP200NTD:ATG13:ATG101 (black) or FIP200NTD:ATG13MR (orange) at 6s time point. Brown represents the overlay of black and orange. Sites of mutation labeled above matching residues. All value are Mean ± SD. (D) Pull-down assays of mutant FIP200 NTD constructs (M1-M9) and wild type with ATG13:ATG101. Both GSH and Strep-Tactin resin were used to pull down GST-FIP200NTD:Strep-ATG13:ATG101 complex from lysate of overexpressing HEK cells. The pull-down results were visualized by SDS-PAGE and Coomassie blue staining. (E) Pull-down assay of mutant FIP200 NTD constructs (4A) and wild type with ATG13:ATG101. (F) Expression level of FIP200 in samples used for mitophagy analysis. (G) Quantification of mito-mKeima ratiometric FACS analysis of WT or FIP200 KO cells re-expressing WT or mutant FIP200 after 5h of OAQ treatment. N=3 biological replicates. All value are Mean ± SD. p value: ** < 0.01; **** < 0.0001; n.s., not significant.

We used HDX-MS to systematically compare FIP200 NTD:ATG13:ATG101 with FIP200 NTD alone to identify regions in FIP200 NTD interacting with ATG13:ATG101 (Fig. 2C, Fig. S2A-I, Table S2). In general, FIP200 NTD had lower H/D exchange when bound to ATG13:ATG101, as the global HDX difference with FIP200 NTD alone was negative (−3.6% for all peptides combined), consistent with the overall stabilization seen in NSEM 2D class averages. Significant protection (< −10 %) was seen in 9 regions of the FIP200 NTD. We mutated side-chains within each region and tested the effects on ATG13:ATG101 binding. Each region was converted a poly Gly-Ser sequence of equal length to the wild-type region being replaced. GST and Strep pulldown assays showed that mutation of regions 1 (73-80), 3 (319-326), 5 (435-442) and 8 (482-489) impaired the stability of FIP200 NTD (Fig. 2D, red, see bottom gel, Fig. S2J) and no conclusion could be drawn as to whether their interaction with ATG13-ATG101 was direct or not. Mutants M2 (158-165), M4 (350-357), M7 (464-471) and M9 (537-544) had no evident loss of FIP200 NTD stability, nor any effect on ATG13:ATG101 pulldown (Fig. 2D, black; top gel, Fig. S2J-K). Thus ATG13:ATG101 binding leads to a large overall decrease in FIP200 NTD dynamics extending across regions beyond those essential for ATG13:ATG101 binding. Mutation of region M6 (443-450) had no loss of protein expression while eliminating the interaction with ATG13:ATG101 (Fig. 2D, bold, Fig. S2J-K). Region 6 is therefore a major locus of ATG13:ATG101 binding.

The 582-585 region that had been previously proposed to be the interaction site for ATG13 (Chen et al., 2016) had a slight decrease in protection, inconsistent with the expectation that the direct interacting regions should show substantial increases in protection. We replicated the 582-585 4A mutant from that study (Chen et al., 2016). ATG13:ATG101 pulled down less FIP200 NTD (4A) than wild type (Fig. 2E), but we attribute this to FIP200 NTD (4A) being expressed at a much lower level than wild-type FIP200 NTD. Taking the observation that the 4A mutant reduces FIP200 NTD stability, together with the lack of an increase in HDX protection in this region upon ATG13 binding, we ascribe the decrease in binding and the phenotype observed by Chen et al., 2016 to decreased stability of the FIP200 NTD dimer, rather than a direct interaction.

Having mapped the interaction sites for ATG13:ATG101 on FIP200, we sought to investigate the function of these sites in an autophagic process. We assessed mitophagy in HeLa cells using a mito-mKeima assay (Vargas et al., 2019). The M6 mutant construct which disrupted the FIP200 binding site for ATG13 interface was transfected in HeLa cells to determine its effects (Fig. 2F-G). FIP200 KO HeLa cells showed a severe defect in mitophagy relative to the wild type HeLa cells (Fig. 2G). Transient transfection of WT FIP200 and FIP200 M6 corresponding to region 6 (443-450 mutated to GS) was carried out as described earlier (Vargas et al., 2019) and mitophagy was assayed. After a 5h OAQ (Oligomycin, Antimycin A and QVD) treatment to induce mitophagy, cells were analyzed using FACS (Fig. S3A). The FIP200 M6 mutant exhibited a 5-fold reduction in mitophagy when compared to WT FIP200 in the KO rescue experiment, confirming that FIP200 binding to ATG13 is important for its function.

### Mapping the FIP200 binding site on ATG13

We used HDX-MS to compare FIP200 NTD:ATG13:ATG101 with ATG13:ATG101 and so identify the regions of ATG13:ATG101 involved in assembly with FIP200 NTD. In general, the ATG13 middle region (363-460) (ATG13 MR) showed reduced HDX (−6.5% for all peptides combined), when bound to the FIP200NTD dimer (Fig. 3A, Table S2). Both the HORMA domain of ATG13 and ATG101 only showed slight differences, below the 10 % threshold of significance (Fig. 3A, Fig. 3B). These data suggest that the ATG13 MR, not the HORMA dimer is the FIP200 NTD binding site. This is consistent with the finding that deletion of ATG13 isoform2 348-373 blocks FIP200 interaction (Wallot-Hieke et al., 2018). This is also consistent with the NSEM result that in the FIP200 NTD:ATG13:ATG101, the HORMA domain dimer density was averaged out in 2D classifications (Fig. 2B).

**Figure 3.**
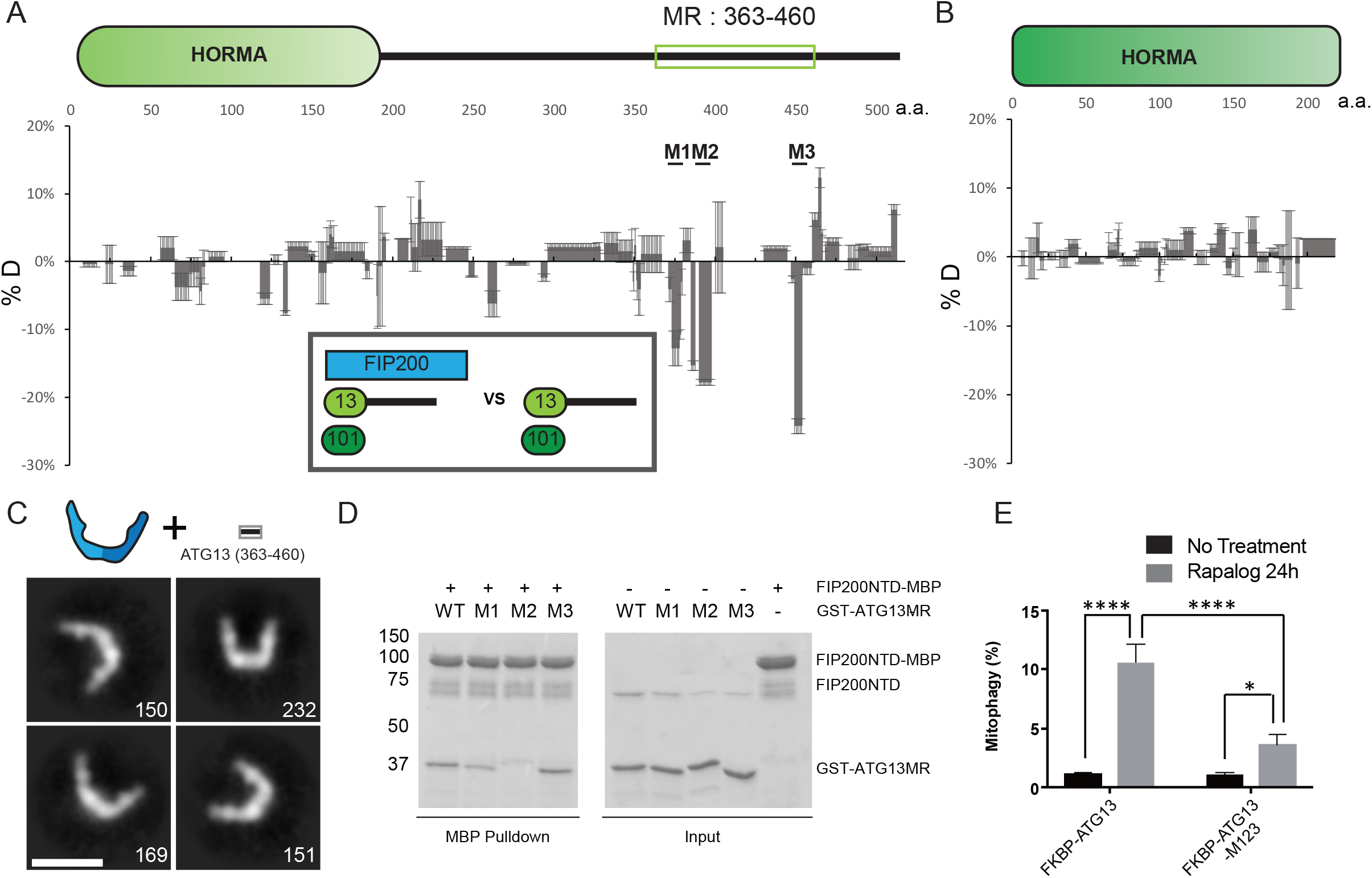
Mapping the FIP200 NTD binding sites on ATG13. (A) Difference of Hydrogen Deuterium Exchange percentages of the ATG13 in ATG13:ATG101 vs ATG13:ATG101:FIP200 at 6s time point. Sites of mutation labeled above matching residues. All value are Mean ± SD. (B) Difference of Hydrogen Deuterium Exchange percentages of the ATG101 in ATG13:ATG101 vs ATG13:ATG101:FIP200. All value are Mean ± SD. (C) Negative stain EM 2D class averages of FIP200 NTD:ATG13 MR complex. Scale bar is 20 nm. (D) Pull-down assays of mutant ATG13MR constructs (M1-M3) and wild type with FIP200NTD. Amylose resin were used to pull down purified GST-ATG13MR:FIP200NTD-MBP complex. The pull-down results were visualized by SDS-PAGE and Coomassie blue staining. (E) Quantification of mito-mKeima ratiometric FACS analysis of HeLa cells stably expressing mito-mKeima-P2A-FRB-Fis1 and FKBP-GFP-ATG13 or mutant after 24 h of Rapalog treatment. N=3 biological replicates. All value are Mean ± SD. p value: * = < 0.05; **** < 0.0001.

To compare the properties of the isolated ATG13 MR with the full ATG13:ATG101 subcomplex, we compared the HDX of FIP200 NTD:ATG13 MR with FIP200 NTD alone (Fig. 2C, Fig. S2A-I, Table S2). The presence of the ATG13 MR led to a pattern of differences identical to those induced by full-length ATG13:ATG101 (Fig. 2C). Furthermore, NSEM analysis of FIP200 NTD:ATG13 MR sample showed stable C shapes like those of the FIP200 NTD:ATG13:ATG101 (Fig. 2B, 3C, Fig S1G, Table S1). Thus, the 98-residue ATG13 MR fully recapitulates the properties of the full ATG13:ATG101 subcomplex with respect to its ability to bind and rigidify FIP200. Three regions in the ATG13 MR (371-378, 390-397 and 446-453) were selected and mutated to poly Gly-Ser sequences of equal length to the wild-type region being replaced. GST tagged ATG13 MR constructs and FIP200 NTD-MBP were purified and used in MBP pulldown assays (Fig. 3D, S2L). Mutation of the region 2 (390-397) largely impaired the interaction between ATG13 MR and FIP200 NTD, while mutation of the region 1 (371-378) moderately impaired the interaction. Slight effect was observed when using mutation of the region 3 (446-453).

It was previously found that in *atg13* KO MEFs rescued by ATG13 construct expression, loss of the N-terminal part of the ATG13 MR has no effect on starvation-induced autophagy and only a slight effect on torin-induced autophagy (Wallot-Hieke et al., 2018). We assessed mitophagy in HeLa cells using a mito-mKeima assay (Vargas et al., 2019). To completely disrupt the FIP200 binding, all three regions in ATG13 MR (371-378, 390-397 and 446-453) were mutated to poly Gly-Ser sequences of equal length to the wild-type region being replaced to generate the M123 mutant. The WT and M123 mutant were then stably expressed at a similar level in HeLa cells and mitophagy was assayed using a previously described chemical-induced dimerization (CID) system (Vargas et al., 2019) (Fig. 3E, Fig. S3B-C). Compared to ATG13 WT, ATG13 M123 mutant triggered a significantly reduced mitophagy response (3-fold reduction). This contrasts with a report that mutating a subset of the FIP200-binding region of ATG13, residues 348-373, does not impact starvation-induced autophagy in MEFs (Wallot-Hieke et al., 2018). These data show that the three FIP200-binding regions of ATG13 analyzed here are functionally important in mitophagy.

### ULK1-EAT interactions with FIP200 NTD-ATG13-ATG101

The FIP200 NTD:ATG13:ATG101:ULK1 complex was purified from HEK 293 cells and subjected to HDX-MS analysis to map the ULK1 binding sites on the rest of the complex. FIP200 NTD peptide 319-326 and ATG13 peptide 482-517 showed significantly decreased HDX in the presence of ULK1 (Fig. 4A, B). In contrast, ATG101 had no significant HDX changes upon the addition of ULK1 (Fig. 4C). The HDX changes in FIP200 and ATG13 peptides suggested that these two regions could serve as the ULK1 binding site. FIP200 region 319-326 is also important for FIP200 NTD stability (M3; Fig. 2D, Fig. S2J). It was reported that the EAT domain of ULK1 interacts with the C terminus of ATG13 (Wallot-Hieke et al., 2018), as anticipated from the homology between the corresponding regions of yeast Atg1 and Atg13 (Fujioka et al., 2014; Stjepanovic et al., 2014). Consistent with this, ULK1 EAT alone induced essentially identical HDX changes in both FIP200 NTD and ATG13 as compared to full-length ULK1 (Fig. 4A, B). To investigate whether ULK1, FIP200 NTD and ATG13 form a three-way interface, ULK1(N-830/ ΔEAT) and ATG13(N-486/ΔC) were assayed by pull-down in the presence of all four ULK1 complex subunits. GST-FIP200 NTD can still pull down some MBP-ULK1 or vice versa even in the presence of ATG13-ΔC (Fig. S4). This demonstrates that once ATG13 brings FIP200 and ULK1 together, a direct interaction exists between FIP200 NTD and ULK1 EAT.

**Figure 4.**
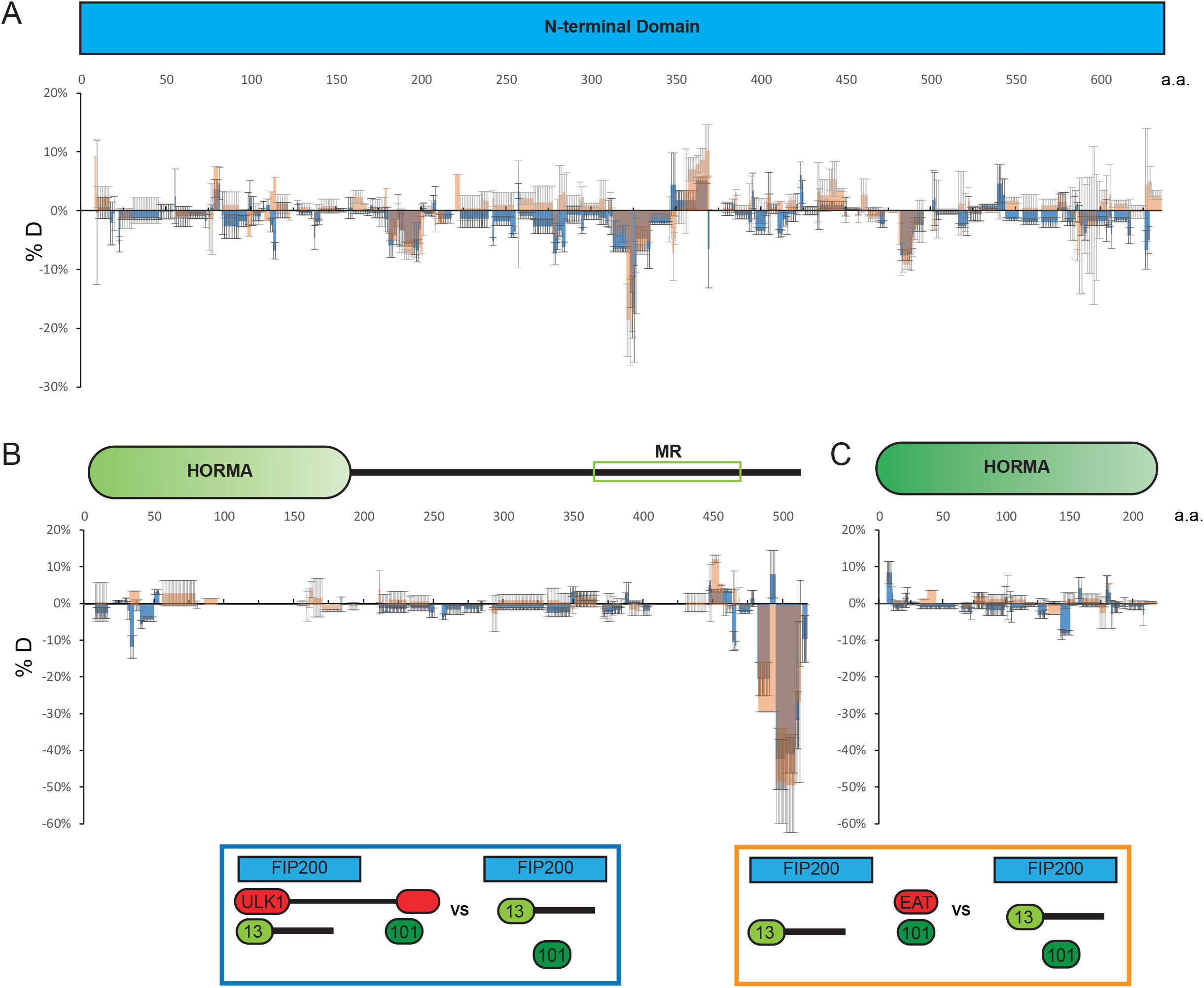
Interactions between ULK1 and the FIP200:ATG13:ATG101 subcomplex. (A-C) Difference of Hydrogen Deuterium Exchange percentages of the FIP200 NTD(A), ATG13(B) and ATG101(C) in ATG13:ATG101:FIP200NTD vs in ATG13:ATG101:FIP200NTD:ULK1(blue) or ATG13:ATG101:FIP200NTD:ULK1 EAT(orange) at the 60s time point. Brown represents the overlay of blue and orange. All value are mean ± SD.

### An asymmetric ULK1 complex with 1:2:1:1 ULK1-FIP200-ATG13-ATG101 stoichiometry

Having defined the minimal interacting regions responsible for assembly of the ULK1 complex, we sought to localize them in space relative to the FIP200 NTD scaffold and to understand their stoichiometry. First, FIP200 NTD:MBP-ATG13 MR was analyzed by NSEM. 2D class averages showed that only one MBP density could be seen near the center of “C”, suggesting that only one molecule of ATG13 binds per FIP200 NTD dimer (Fig. 5A, Fig. S1H, Table S1). This suggests that the ATG13 binding site spans both FIP200 NTD monomers in the C shaped dimer.

**Figure 5.**
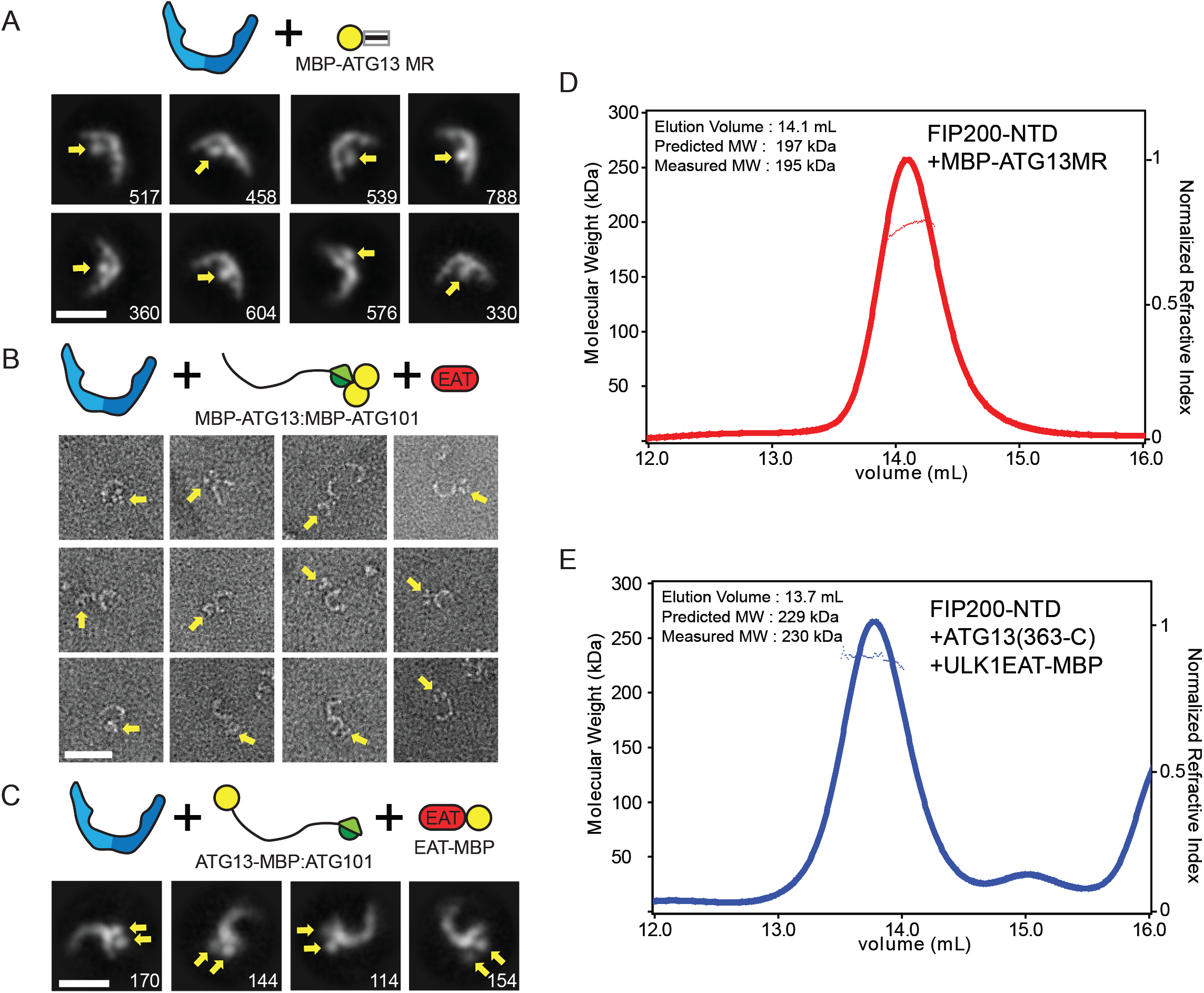
The ULK complex has unequal subunit stoichiometry. (A) Negative stain 2D class averages of FIP200 NTD:MBP-ATG13 MR complex. Densities corresponding to MBP tags are labeled with yellow arrows. Scale bar is 20 nm. (B) Single particles of FIP200 NTD:MBP-ATG13:MBP-ATG101:ULK1 EAT complex. Density for dual MBP tagged HORMA domains highlight with yellow arrows. Scale bar is 20 nm. (C) 2D class averages of FIP200 NTD:ATG13-MBP:ATG101:ULK1 EAT-MBP complex. Densities corresponding to MBP tags are labeled with yellow arrows. Scale bar is 20 nm. (D-E) Multiangle light scattering traces for FIP200 NTD:MBP-ATG13MR (D) and FIP200 NTD:ATG13(363-C):ULK1 EAT-MBP (E).

To determine if the ATG13-ATG101 HORMA dimer was structurally ordered with respect to the FIP200 NTD scaffold, we fused MBP tags to each protein. This more than doubles the effective mass of these domains, making them visible by NSEM. In single particles of FIP200 NTD:MBP-ATG13:MBP-ATG101:ULK1 EAT a trimer density can be seen in the vicinity of the FIP200 NTD dimer, corresponding to density for the HORMA dimer and the two MBP tags present (Figure 5B, Fig. S1I, Table S1). 2D classification of the complex showed a stable FIP200 NTD dimer but most averages had no additional densities for the MBP tags (Fig. S1I). We interpret this to mean that the HORMA dimer does not directly interact with, and is not ordered with respect to, FIP200 NTD.

We next carried out NSEM with ATG13-MBP and ULK1 EAT-MBP. MBP tags placed at the ATG13 C-terminus and on the ULK1 EAT domain were co-localized as seen by the presence of two extra density lobes present at the tip of one arm of the FIP200 NTD dimer (Fig. 5C, Fig. S1J, Table S1). 2D classification showed that the ULK1 EAT and the C terminus of ATG13 are located near one of the tips of the FIP200 NTD “C” shape.

The results of the MBP tagging experiments implied that the ULK1 complex is asymmetric and has nonequal subunit stoichiometry. To test this, we used MALS to determine the stoichiometry of the ULK1 complex by direct determination of the molecular mass. FIP200 NTD:MBP-ATG13 MR and FIP200 NTD:ATG13(363-C):ULK1 EAT-MBP both showed molecular weights consist with a stoichiometry of two molecules of FIP200 NTD for each one molecule of all other components (Fig. 5D, E). The measured molecular weights of 195 kDa and 230 kDa correspond to the expected mass of complexes with 2:1 ratios between the FIP200 NTD and the other subunits (197 kDa, and 229 kDa, respectively). Taken together, the NSEM and MALS data show that the FIP200 NTD dimer assembles asymmetrically with one copy each of ATG13, ATG101 and ULK1.

### Cryo-EM structure of ATG13 MR-bound FIP200 NTD

We used cryo-EM to investigate the architecture of the FIP200 NTD dimer:ATG13 MR complex at higher resolution. The narrow extended C-shape and flexible nature of FIP200 made it a challenging sample for both sample preparation and data collection. Nevertheless, the use of several technical improvements made it possible to obtain an intermediate resolution structure. Graphene oxide was used to protect the FIP200 NTD dimer from the air water interface and allowed collection of high contrast micrographs (Fig. S5A). Accurate particle picking was critical for the centering of FIP200 NTD C-shape. The neural network based crYOLO picker (Wagner et al., 2019) was trained by manual picking and used to autopick micrographs from three datasets (Fig. S5B, Table S3). 2D classification showed the FIP200 NTD dimer in an array of more or less open conformational states similar to that seen in the NSEM data (Figure 6A). A distinct bump was observed on the inner rim of the dimer. After 3D classification full dimer maps were resolved between 12-15Å resolution (Figure 6B, Fig. S5B). When overlaid these maps show a large range of tip-to-tip distances, spanning 160Å to 220 Å across.

**Figure 6.**
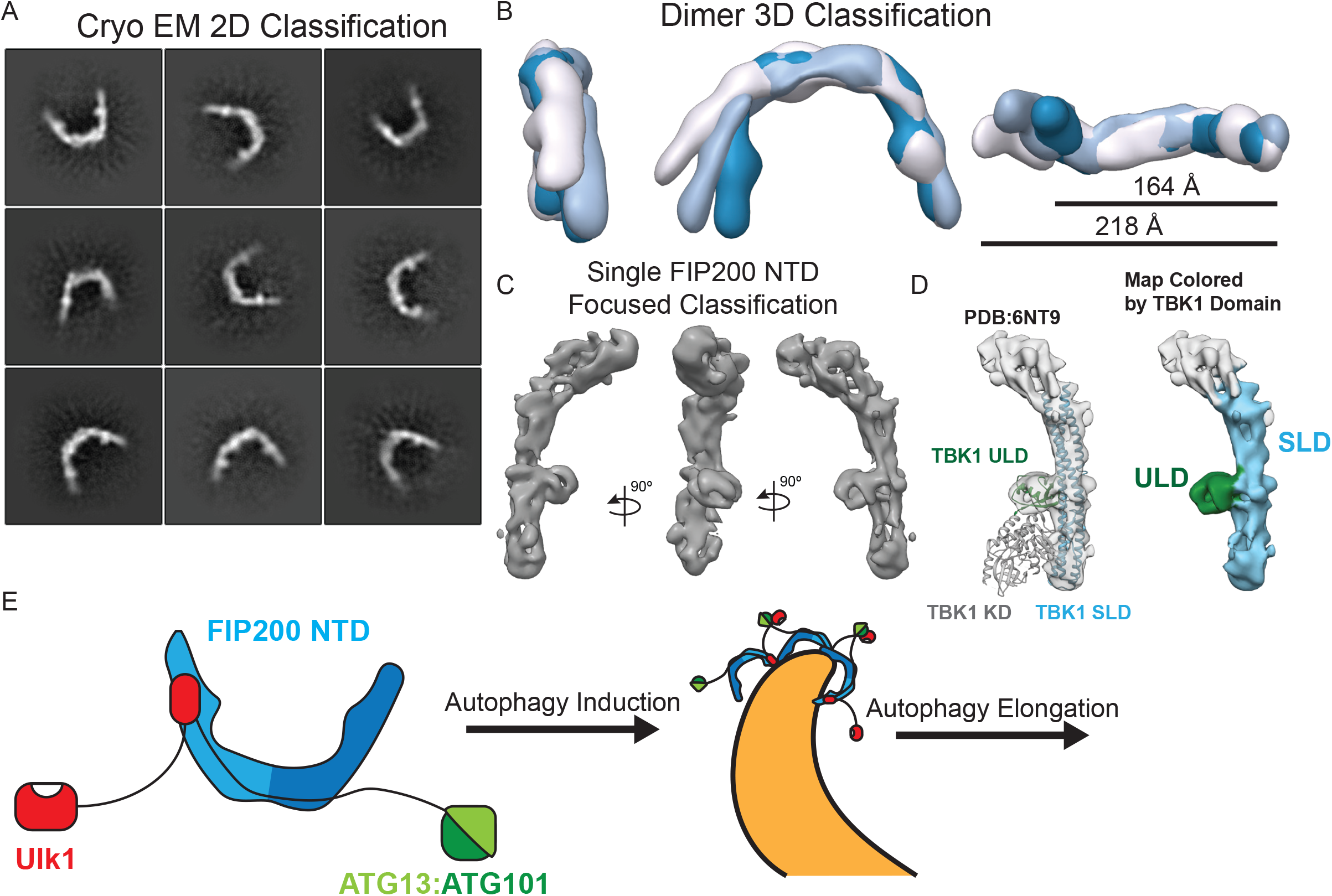
Cryo EM reconstruction of the FIP200 NTD:ATG13 MR complex and model for ULK complex assembly. (A) CryoEM 2D class averages of FIP200 NTD:ATG13 MR. (B) 3D reconstructions of dimeric FIP200 NTD overlaid with measurements from tip-to-tip ranging from 164Å to 218Å across. (C) Views of masked final 3D reconstruction of FIP200 NTD monomer. (D) Fit of TBK1 structure into the final map (left) and colored by ubiquitin like domain (green) and scaffold like domain (blue) (right) (E) Model for asymmetric FIP200 ULK complex assembly and proposed formation of higher over oligomers during autophagy initiation.

To improve the resolution of the FIP200 NTD we used masked focused refinement on a single arm improved the resolution to ~9Å (Figure 6C, Fig. S5B-E). At this resolution helical densities can be seen which run along the length of the C shape. Additionally, more features are seen at density within the dimer C shape.

To interpret this moderate resolution map, given the lack of pre-existing atomic structures for FIP200 outside of the CLAW domain, we relied on structure predictions servers. Robetta analysis (Song et al., 2013) suggested that the TBK1 scaffold like (SLD) and ubiquitin like (ULD) domains (Shu et al., 2013) as the basis for a structural model. Docking of TBK1 ULD and SLD (Shu et al., 2013; Zhang et al., 2019) showed a good fit into our density (Fig. 6D left) while another candidate, the yeast Atg17 structure, did not (Fig. S5F). These domains from TBK1 are comprised of 429 residues which would not account for the FIP200 NTD construct (1-640) (Fig 6D right). The dimer interface of the FIP200 NTD dimer remains unmodeled at our current resolution. The bump seen in 2D averages corresponds to the ULD and the backbone of the arm forms a helical bundle which is similar in spacing and structure to the SLD of TBK1 (Figure 6D). TBK1 has a conserved interface positioned between the ULD domain and the scaffolding domain containing residues NPIF (376-380) (Shu et al., 2013; Zhang et al., 2019). FIP200 contains the same sequence near the end of the predicted ULD in residues 67-72 (Fig. S5G). This region corresponds with a region of conserved primary sequence between FIP200 and TBK1.

## Discussion

The ULK1 complex has a central role in autophagy initiation, yet its structural organization (Lin and Hurley, 2016) is not nearly as well understood as that of the other major complexes of autophagy initiation, PI3KC3-C1 and ATG16L1-ATG5-ATG12. In particular, there have been no structural data for the largest subunit, FIP200, apart from very recent structures of the Claw domain (Turco et al., 2019), representing less than 10% of the total mass of the protein. Here we have substantially filled these gap at two levels. We have mapped the overall structural organization and showed that the FIP200 NTD is the hub about which the larger complex is organized. We infer that the FIP200 CC and Claw, the ATG13-ATG101 HORMA dimer, and the ULK1 kinase domain probably project away from the hub, given that they are non-essential for complex formation, do not show HDX protection patterns consistent with interactions, or both (Fig. 6E). These three projecting regions appear not to interact with the hub at all under the conditions of these studies.

FIP200 is often considered to be the functional ortholog of Atg17 in mammals (Lin and Hurley, 2016; Mizushima, 2010). Like Atg17, FIP200 NTD dimerizes through its C-terminus, and like Atg17, the N-terminus is located near the outer tips of the subunits in the dimer context. On the other hand, the FIP200 dimer gives rise to a C shape, in contrast to the S-shape of Atg17 (Ragusa et al., 2012). The Atg17 S has been postulated to have a unique role in nucleating cup-shaped phagophores in bulk autophagy (Bahrami et al., 2017) on the basis of first-principles physical membrane modeling, something that would not be achieved by a C. Cup-shaped FIP200 structures have been visualized in amino acid-starved U2OS cells by super-resolution microscopy (Kenny, 2019), likely corresponding to phagophores observed in cell by EM (Hayashi-Nishino et al., 2009; Yla-Anttila et al., 2009). The ~20 nm diameter of the FIP200 C shape is potentially compatible with localization on highly curved ATG9 vesicles or on the rim of the growing phagophore.

We mapped the mutual determinants for FIP200 and ATG13 binding to one another. Given that each of these proteins is central to autophagy, that they interact strongly with one another, and that the yeast Atg13-Atg17 interaction is functionally critical, we had expected that weakening this interaction would lead to a reduction in function. We confirmed that mutating the binding site for ATG13 on FIP200, and the FIP200 binding site of ATG13, impaired mitophagy, consistent with expectation.

The similarity in structure between the FIP200 NTD and the combined ULD and scaffold domains of TBK1 (Shu et al., 2013; Zhang et al., 2019) was unexpected. Despite the limited resolution of the NTD cryo-EM structure, the positions of the helices and ULD relative to one another are unmistakably related and otherwise unique. This suggests to us that FIP200 is a structural composite of a TBK1-like N-terminal region and an Atg11-like C-terminal region. TBK1 in turn appears to be a structural chimera of a kinase with the FIP200 NTD. Given increasing evidence for ubiquitous roles for TBK1 in mammalian autophagy initiation (Vargas et al., 2019) it should perhaps not be so surprising that the FIP200 and ULK1 system, collectively, and TBK1 have a structural and, presumably, evolutionary relationship.

We carried out an intensive examination of the network of interactions between the ULK1 complex subunits, and came to one more unexpected conclusion, that the complex contains a constitutive dimer of FIP200 but only one copy of every other subunit. The C-terminal end of FIP200 NTD appears to be close to the center of the dimer interface, thus the start of the dimeric coiled coil domain projects away from this interface with matching symmetry. The division of labor of FIP200 domains seems to be that the Claw dimer at the very tip of the complex binds cargo, the coiled-coil acts as a long-range connector, and the NTD coordinates the ATG13-ATG101 subcomplex and ULK1 itself. As a stable dimer even at high dilution in our experiments, it seems unlikely that FIP200 requires cargo to dimerize. Therefore, a receptor-like model in which FIP200 transmitted a signal by undergoing cargo-induced dimerization seems unlikely to us. The 2:1 FIP200:ULK1 stoichiometry would prevent ULK1 from dimerization and auto-activating in the absence of higher-order clustering. This suggests an appealing mechanism for cargo-induced ULK1 activation in which multiple FIP200 dimers cluster on the autophagic substrate/cargo, bringing ULK1 monomers into proximity for autoactivation.

## Acknowledgements

We thank P. Grob and D. Toso for cryo-EM facility support. This work was supported by the NIH (R01 GM111730 to J.H.H. and F99 CA223029 to L.N.Y.), HFSP (RGP0026/2017 to J.H.H.), and the Jane Coffin Childs Foundation (A. L. Y.). This research was supported (in part) by the Intramural Research Program of the NIH, NINDS. J.H.H. is a scientific founder of and receives research funding from Casma Therapeutics.

## Methods

### Plasmid construction

The sequence of all DNAs encoding components of ULK1 complex were codon optimized, synthesized and then subcloned into the pCAG vector. All components were tagged with GST, MBP or TwinStrep-Flag(TSF) for affinity purification or pull down assays. N-terminal GST, MBP or TSF tags may be followed by a tobacco etch virus cleavage site (TEVcs). All constructs were verified by DNA sequencing. Details are shown in Table S4.

### Protein expression and purification

All proteins used for NSEM, HDX-MS and MALS analysis were expressed in HEK293-GnT1 suspension cells by using the polyethylenimine (Polysciences) transfection system. Cells were transfected at a concentration of 2-2.5 × 10^6^/mL and harvested after 48 hours. The harvested cells were pelleted at 500 × g for 5 min at 4°C, and then washed with PBS once. The pellets were then lysed with lysis buffer containing 50 mM Tris-HCl pH 7.4, 200 mM NaCl, 2 mM MgCl2, 1 mM TCEP, 1% Triton X-100, 10% Glycerol and protease inhibitors (Roche) before being cleared at 16000 × g for 30 min at 4°C. The supernatant was then incubated with Glutathione Sepharose 4B (GE Healthcare), Amylose resin (New England Biolabs) or Strep-Tactin Sepharose (IBA Lifesciences) as appropriate, with gentle shaking for 12 hours at 4°C. The mixture was then loaded onto a gravity flow column, and the resin was washed extensively with wash buffer (50mM HEPES pH8.0, 200mM NaCl, 1mM MgCl2 and 1mM TCEP). The proteins were eluted with wash buffer containing 50 mM glutathione, 50 mM maltose or 10 mM desthiobiotin, as appropriate. In some cases, two affinity steps were used. Constructs containing TEV cleavage sites were treated with TEV protease at 4°C overnight. For HDX-MS and NSEM analysis, the protein was applied to a final size exclusion chromatography step before use. For the Strep-ATG13:Strep-ATG101 complex, a Superdex 200 column (GE Healthcare) was used, and for all other samples, a Superose 6 column (GE Healthcare) was used.

### Pull-down assays

10 mL of HEK293-GnT1 suspension cells were transfected at the concentration of 2-2.5 × 10^6^/mL and harvested after 48 hours. The harvested cells were pelleted at 500 × g for 5 min at 4°C, and then washed with 5mL PBS once. The pellets were then lysed with 1mL lysis buffer containing 50 mM Tris-HCl pH 7.4, 200 mM NaCl, 2 mM MgCl_2_, 1 mM TCEP, 1% Triton X-100, 10% Glycerol and protease inhibitors (Roche) before being cleared at 12000 rpm for 10 min at 4°C. The supernatant was then incubated with 20 μL Glutathione Sepharose 4B (GE Healthcare), Amylose resin (New England Biolabs) or Strep-Tactin Sepharose (IBA Lifesciences) with gentle shaking for 8 hours at 4°C. The protein-bound resin was washed with 1 mL lysis buffer 3 times, and then eluted with 60 μl elution buffer containing 50 mM glutathione, 50 mM maltose or 10mM desthiobiotin, respectively. The eluted proteins were applied to SDS–PAGE for analysis. For Fig. 1B, FIP200(N-640)-MBP and FIP200(636-C)-MBP were first purified by GST affinity purification, followed by TEV cleavage and then MBP affinity purification. The MBP-ULK1:Strep-ATG13:Strep-ATG101 subcomplex was purified by Strep affinity purification, and then left on the resin. The resin was then mixed with FIP200 protein (final concentration: 200nM) at 4°C with gentle shaking. For Fig. 2A, purified GST-FIP200, MBP-Strep-ULK1 and Strep-ATG13:Strep-ATG101 were used. The final buffer was 20mM HEPES pH8.0, 200 mM NaCl, 1 mM TCEP, 5 mM desthiobiotin and 1% Triton-X-100. The protein concentration was 200 nM. For Fig. 3D, purified FIP200-MBP and GST-ATG13MR were used. The final buffer was 20mM HEPES pH8.0, 200 mM NaCl, 1 mM TCEP and 1% Triton-X-100. The protein concentration was 500 nM.

For quantification of the pulldown assay in Fig. 2D and 3D, ImageJ was invited to measure the intensity of bands in SDS-PAGE gel. The expression of GST-FIP200NTD was calculated as Intensity(GST-FIP200NTD in GST pull-down gel) / Intensity(Strep-ATG13 in Strep pull-down gel). Relative expression efficiency of GST-FIP200NTD was calculated as Expression(GST-FIP200 WT or Mutants) / Expression(GST-FIP200 WT). The pulldown of GST-FIP200NTD by Strep-ATG13:ATG101 was calculated as Intensity(GST-FIP200NTD in Strep pull-down gel) / Intensity(GST-FIP200NTD in GST pull-down gel) / Intensity(Strep-ATG13 in Strep pull-down gel). Relative pulldown efficiency of GST-FIP200NTD was calculated as Pulldown(GST-FIP200 WT or Mutants) / Pulldown(GST-FIP200 WT). The pulldown of GST-ATG13MR by FIP200NTD-MBP was calculated as Intensity(GST-ATG13MR in MBP pull-down gel) / Intensity(FIP200NTD-MBP in MBP pull-down gel) / Intensity(GST-ATG13MR in Input gel). Relative pull-down efficiency of GST-ATG13MR was calculated as Pulldown(GST-ATG13MR WT or Mutants) / Pulldown(GST-ATG13MR).

### Hydrogen-deuterium exchange mass spectrometry

Protein samples for HDX were concentrated to a 10 μM stock before H/D exchange. The exchange was initiated by adding 95 μL of deuterated buffer containing 20 mM HEPES (pH 8.0), 200 mM NaCl, 1 mM TCEP into 5 μL of protein stock at 30°C. Exchange was carried out for 6 s, 60 s, 600 s, or 60000 s, and quenched at 0°C by the addition 100 μL of ice-cold quench buffer (400 mM KH_2_PO_4_/H_3_PO_4_, pH 2.2). The 60000 s sample served as the maximally labeled control. All HDX reactions were repeated three times. Quenched samples were injected into a chilled HPLC (Agilent) setup with in-line peptic digestion and desalting steps. The analytical column used was a Biobasic 8.5 μm KAPPA column (Fisher Scientific). The peptides were eluted with an acetonitrile gradient and electrosprayed into an Orbitrap Discovery mass spectrometer (Thermo Scientific) for analysis. To generate the gradient, solvent A was 0.05% TFA, while solvent B was 0.05% TFA in 90% acetonitrile. The elution method was as follows: 0-6min: 10%B; 6-18min: from 10%B to 55%B; 18-19min: from 55%B to 90%B; 19-25min: 90%B; 25-26min: from 90%B to 10%B; 26-30min: 10%B. The spray voltage was set at 3.4 kV, capillary temperature was set at 275°C, capillary voltage was set at 37V and tube-lens was set at 120V. As a control, unexchanged samples went through the same process, except that deuterated buffer was replaced by undeuterated buffer.

To identify peptides, unexchanged samples were analyzed by tandem MS/MS analysis with the same HPLC method. Tandem MS/MS was performed using data dependent analysis, in which a cycle of one full-scan MS spectrum (m/z 300-2000) was acquired followed by MS/MS events (CID fragmentation). MS/MS was sequentially generated on the first to the ten most intense ions selected from the full MS spectrum at a 35% normalized collision energy. The ion trap analyzer was used for MS2, activation time was 30ms and the dynamic exclusion was set at 30 sec. For HDX mass analysis, only a full-scan MS spectrum was acquired, and the resolution was 30000.

Database searches were performed with the Proteome Discoverer 2.1 (Thermo Fisher Scientific) using the Sequest HT search engine to identify peptides. Raw data were searched against the small database containing all four components of ULK1 complex. The following search parameters were used: unspecific cleavage was used; precursor mass tolerance was set to ± 10 ppm and fragment mass tolerance was set to ± 0.6 Da. Target FDR was set to 1% as the filter cut-off for the identified peptides. For HDX analysis, mass analysis of the peptide centroids was performed using HDExaminer (Sierra Analytics), followed by manual verification for every peptide.

### Negative Stain Electron Microscopy Collection and Processing

Purified protein samples were diluted to ~50-200 nM final concentration in running buffer immediately before application to glow discharged continuous carbon grids. Protein samples were stained twice with 2-4% uranyl acetate and allowed to dry at room temperature. Samples were imaged using a T12 or F20 transmission electron microscope operating at 120 keV (Thermo Fisher) as indicated in Table S1. Data were manually collected and assessed for stain quality throughout data collection. F20 datasets used a recorded magnification of 89,000X, collected with a ultrascan camera at a pixel size of 1.5 Å/pixel (Gatan). T12 datasets were captured using a magnification of 49,000X with a 4k × 4k CCD camera (Gatan) which corresponds to 2.2 Å/pixel.

All datasets spanned a range of 1-4 μm defocus and a total dose of 30-50 e-/Å^2^. Single particles were selected using Relion Autopicker (Zivanov et al., 2018) and extracted with the indicated box size (Table S1). Data were cleaned by 2D classification within Cryosparc2, removing classes which had no features or contained background picks. Final 2D classification into 80 or 50 classes was performed with the ‘uncertainty parameter’ set to 8. This setting yielded the best distribution of class averages across all datasets.

### Size Exclusion and Multi-angle Light Scattering Analysis

All light scattering experiments were performed with a running buffer of 50 mM Tris, 150 mM NaCl, 1 mM TCEP at pH 7.8. Purified ULK complex sample was concentrated to ~6-10 μM and 100 μL was injected over a 24 mL Superose 6 Increase 10/300 GL column (GE) in tandem with light scattering analysis using both an Optilab rEX differential refractive index and a DAWN HELEOS II MALS detectors. Data were analyzed using ASTRA VI software (Wyatt Technology) with peak alignments, normalization and band broadening effects determined from a standard of 2 mg/mL bovine serum albumin. The molecular weight of the sample was reported by selecting the peak with the largest UV intensity and averaging the molar mass values across the width of the peak. The radius of hydration was inconsistent across different runs and was not included in the analysis.

### Mitophagy assay

For chemical-induced dimerization (CID) assay, HeLa cells stably expressing mKeima-P2A-FRB-Fis1 and FKBP-GFP-ATG13 and mutants were treated with A/C heterodimerizer (for simplicity, it is called “rapalog” in the figure) (Clontech#635056) for 24 h and then subjected to FACS analysis as previously described (Vargas et al., 2019). For FIP200 KO rescue experiments, HeLa or FIP200 KO cells stably expressing mito-mKeima were co-transfected with 0.25 μg pEYFP-Parkin, 1 μg pHAGE-HA-FIP200-IRES-puro and mutants with FuGENE HD (Promega) for 18 hrs and then treated with 10 μM Oligomycin (Calbiochem), 10 μM Antimycin A (Sigma) and 20 μM QVD (ApexBio) for 5 h before FACS analysis. All the constructs (including site mutagenesis) were made with Gibson assembly (NEB#E2611S) and confirmed by Sanger sequencing. The complete sequence map of each construct is available upon request.

### Cryo Electron Microscopy Sample Preparation and Data Collection

UltrAUfoil 2/2 300 mesh gold grids (Quantifoil) were used for open hole data collection (Dataset 1). Samples were concentrated after gel filtration to ~5 uM and applied to glow discharged grids. Blotting was performed at 100% humidity in a Vitrobot Mark IV (Thermo Fischer) for 2-6 seconds. Graphene oxide coated grid datasets were prepared as follows. UltrAUfoil 1.2/1.3 300 mesh gold grids (Quantifoil) were glow discharged under mild conditions i.e. 10 mAmp for 15 sec. Grids were incubated with a layer of PEI at 1 mg/mL for 2 minutes after which the solution was blotted off using Whatman 1 filter paper followed by two rounds of washing with water. Grids were allowed to dry for 15 minutes and then incubated with 4 μL of graphene oxide flakes (Sigma) at ~0.2mg/mL for 1-2 minutes. Excess graphene oxide solution was wicked away and washed with two 4 μL drops of water. Coated grids were screened in a T12 microscope to assess coverage and quality of graphene oxide before plunge freezing. FIP200:ATG13MR samples were checked via NSEM directly prior to freezing to assess quality of protein and protein concentration. Protein samples for the graphene oxide datasets were diluted immediately before freezing in gel filtration buffer (50 mM HEPES pH 8.0, 200 mM NaCl, 1 mM MgCl_2_ and 1 mM TCEP) to a final concentration between 200-500 nM. 3.0 μL of sample was loaded onto Graphene Oxide coated gold grids. Sample was plunge-frozen using a vitrobot Mark IV (Thermo Fischer) with blot force 10-20 for 3-7 seconds.

Data were collected by the same procedure for all data sets. Samples were clipped and loaded into a Talos Arctica (Thermo Fischer) operating at 200 kV. Frames were collected at 36,000X nominal magnification on a K3 direct electron detector (Gatan) in super-resolution counted mode at 0.5685 Å/pixel. Serial EM was used for automated image shift data collection of a five-target cross pattern. Movies were taken in 100 ms frames at ~1 e^−^/frame, totaling an electron dose of 60 electrons per movie.

### Cryo Electron Microscopy Processing

The data processing scheme for the final maps is shown in Fig. S5. Micrographs were drift corrected using MotionCor2 (Zheng et al., 2017) and Fourier binned to 1.137Å/pixel. CTFFIND4 (Rohou and Grigorieff, 2015) was used to estimation the CTF parameters of the integrated micrographs. Micrographs were cleaned by inspection and FFT quality.

The three datasets were collected and combined during processing as follows. For Datasets 1 and 2 particles were picked using the Relion autopicker and extracted at a box size of 200×200 pixels with 2.274 Å/pixel. Dataset 1 was taken in open holes on a MBP-FIP200 NTD:ATG13 (363-460) sample. Nearly half of the micrographs were removed due to poor drift correction, imaging of empty holes or distorted CTF information. Relion autopicker yielded ~293k initial particle picks for this dataset. Single particles were pruned by 2D classification within Cryosparc2 (Punjani et al., 2017). Although extensive processing schemes were attempted no high-quality 3D initial models were found. This we suspect is due to the shape of FIP200 and the low contrast of the protein in vitreous ice.

Dataset 2 was taken on graphene oxide with FIP200 NTD: ATG13 (363-460) as the protein sample. Micrographs suffering from large graphene oxide creases were removed during early processing steps. Similar particle picking and pruning was performed on this dataset and ~200k initial particle picks were found. Combination of these datasets led to our final reconstruction via a soft mask around the highest density arm of the FIP200 dimer as shown in Fig. S5B. The final 29,198 particles were masked and processed using nonuniform refinement in Cryosparc2. Preferred orientation was seen throughout the steps of processing and was present in both orientation plot and 3DFSC (28671674) of the final single arm map. The final reconstruction was locally filtered and sharpened with a b factor of −661 within Cryosparc2. Of note, multibody refinement, local motion correction, symmetry expansion and additionally 3D classification did not improve the resolution of the final map. Processing within other software packages yielded maps of worse quality and higher degrees of anisotropic density.

Dataset 3 was taken to increase the initial particle count for the graphene oxide data. A similar number of micrographs containing graphene oxide creases was present and removed as compared to Dataset 2. Dataset 3 was picked using a trained model within crYOLO (Wagner et al., 2019) which proved to center on the FIP200 dimer better than other particle pickers in our hands. 2D classification was used to prune Dataset 3 down to ~80k final particles from an initial ~159k particles (Fig. 6A). Upon the realization that crYOLO was performing better for picking, Dataset 1 and 2 were reprocessed leading to final particle counts of ~65k and ~69k particles, respectively. The final map was deposited into the EMDB under accession code 21325.

Combination of the three datasets did not yield a better final resolution model but it did reveal a large conformational landscape of the FIP200 dimer (Fig. 6B). Conformations spanned a 60Å range from ~160-220Å. This information helped to explain our unsuccessful attempts at multibody refinement and local symmetry expansion as motion at this scale has not been resolved that we are aware of. Data were processed together in Cryosparc2 and pruned to a final heterogeneous classification of 213,156 particles. 3D classification into 6 classes yielded the final 3 maps which show strong density for each arm of the FIP200 dimer. Similar to processing of the two dataset final map further processing did not yield higher resolution features.

For docking of the TBK1 and ATG17 structures into our final monomeric density map, we used the “Fit_in_Map” function within Chimera. For the TBK1 ULD/SDD structure we removed the atomic coordinates of the Kinase domain before docking. The structure was first docked manually near the FIP200 NTD monomer and then 1000 fits were sampled with a search radius of 2. The docking position with the highest CC is shown Fig. 6D. Similarly, ATG17 monomer was docked in the same manner and the highest docked position is shown Fig. S5F.

---

**Figure S1.**
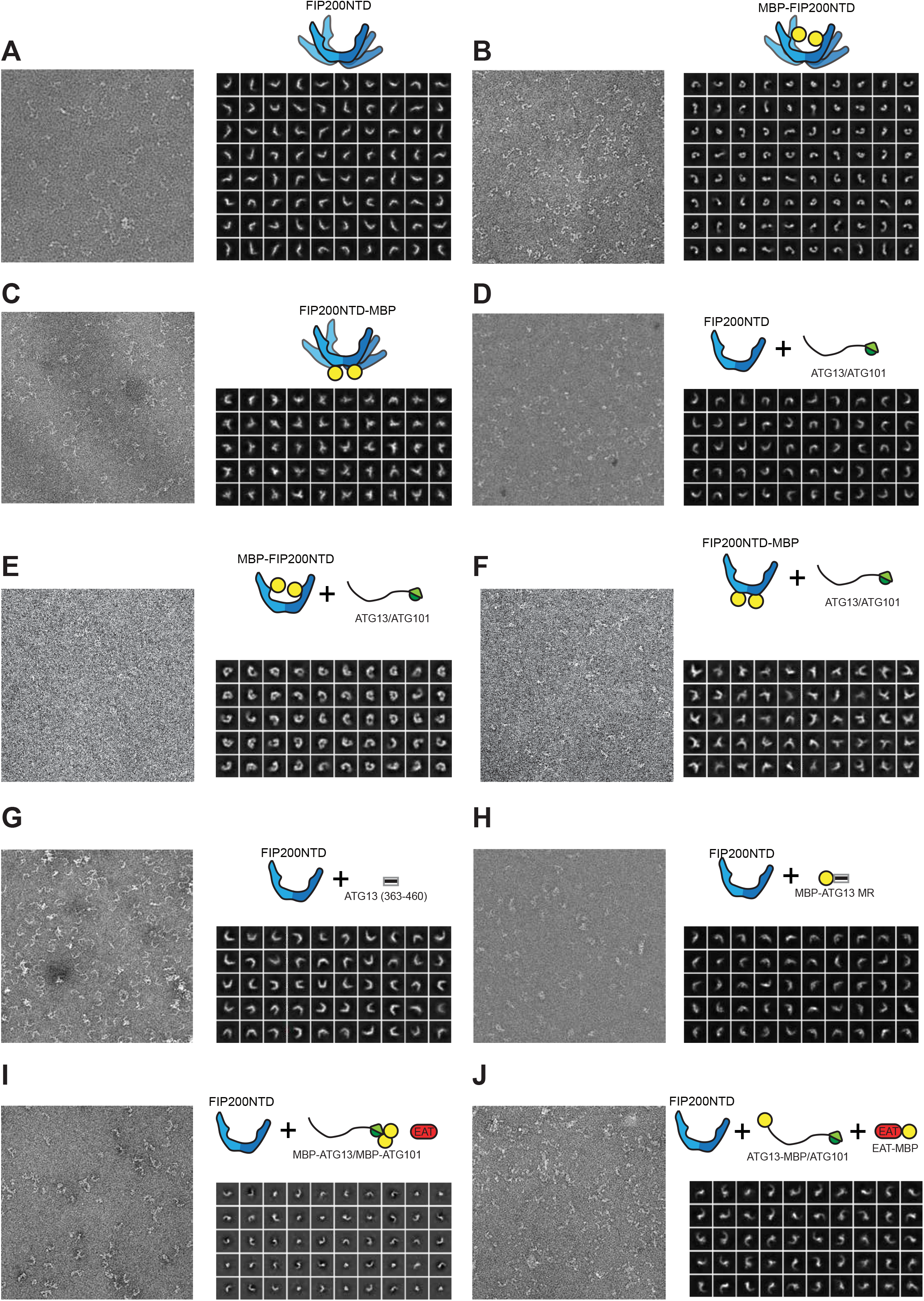
Negative Stain electron microscopy of FIP200 NTD complexes. (A-J) Representative micrograph, schematic and gallery of 2D class averages for each negative stain dataset.

**Figure S2.**
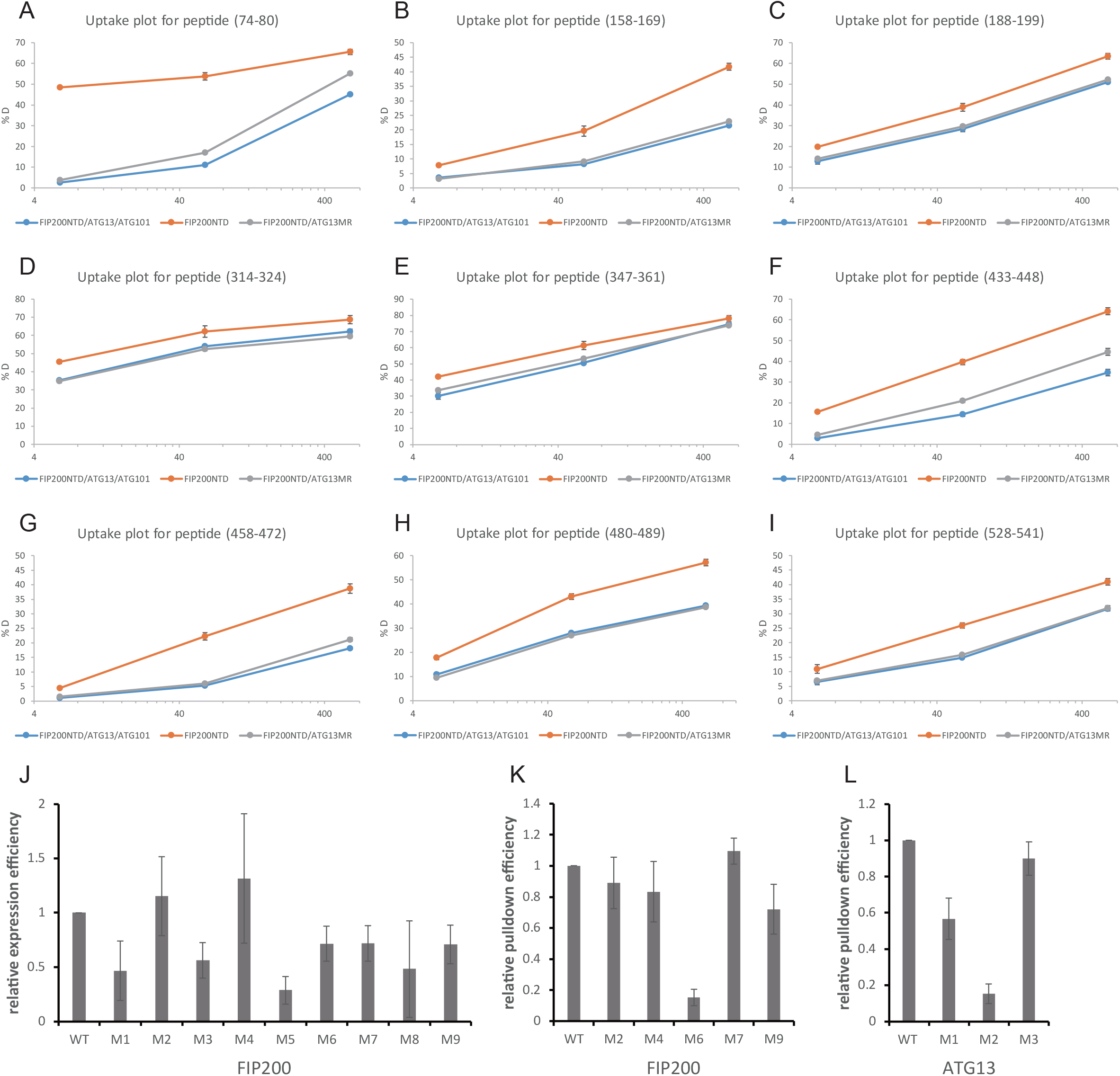
Deuterium uptake plot for peptides in FIP200NTD with significant differences and quantification of pulldown assay. (A) Deuterium uptake plot for peptide (74-80). (B) Deuterium uptake plot for peptide (158-169). (C) Deuterium uptake plot for peptide (188-199). (D) Deuterium uptake plot for peptide (314-324). (E) Deuterium uptake plot for peptide (347-361). (F) Deuterium uptake plot for peptide (433-448). (G) Deuterium uptake plot for peptide (458-472). (H) Deuterium uptake plot for peptide (480-489). (I) Deuterium uptake plot for peptide (528-541). (J) Quantification of relative expression efficiency for GST-FIP200NTD in Fig. 2D. (K) Quantification of relative pull-down efficiency for GST-FIP200NTD in Fig. 2D. (L) Quantification of relative pull-down efficiency for GST-ATG13MR in Fig. 3D. All values are Mean ± SD.

**Figure S3.**
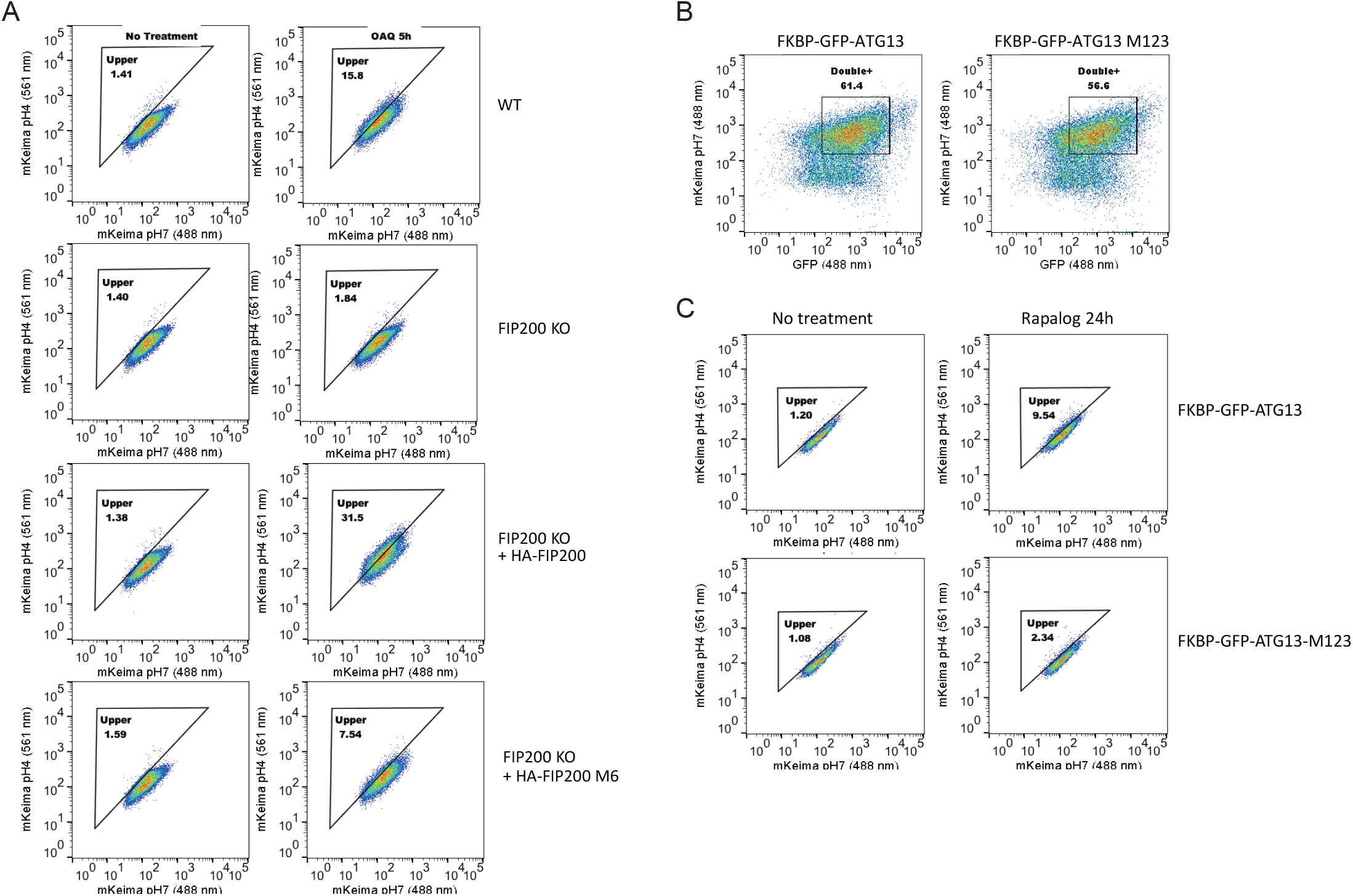
FACS analysis of the FIP200-ATG13 interaction in mitophagy. (A) FACS plots showing mito-mKeima ratio (561/488 nm) for Figure 2G. (B) The WT ATG13 and ATG13 M123 are expressed at similar levels in cells used in Figure 3E. (C) FACS plots showing mito-mKeima ratio (561/488 nm) for Figure 3E.

**Figure S4.**
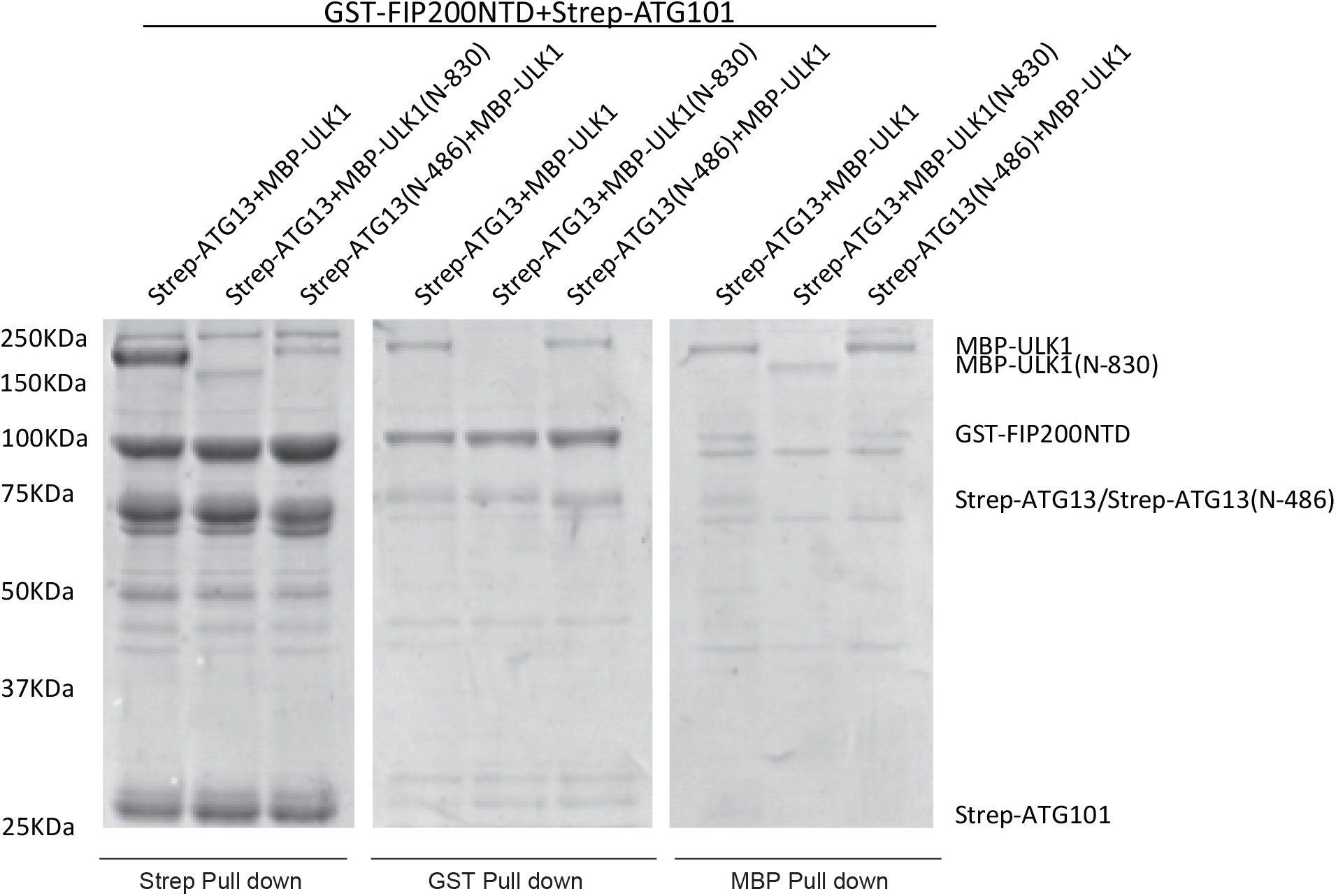
Pull-down assays of ULK1 complex with FIP200 NTD, ATG13 and ULK1 constructs. GSH, Strep-Tactin and Amylose resin were used to pull down GST-FIP200NTD:Strep-ATG13:Strep-ATG101:MBP-ULK1 complex from lysate of overexpressing HEK cells. The pull-down results were visualized by SDS-PAGE and Coomassie blue staining.

**Figure S5.**
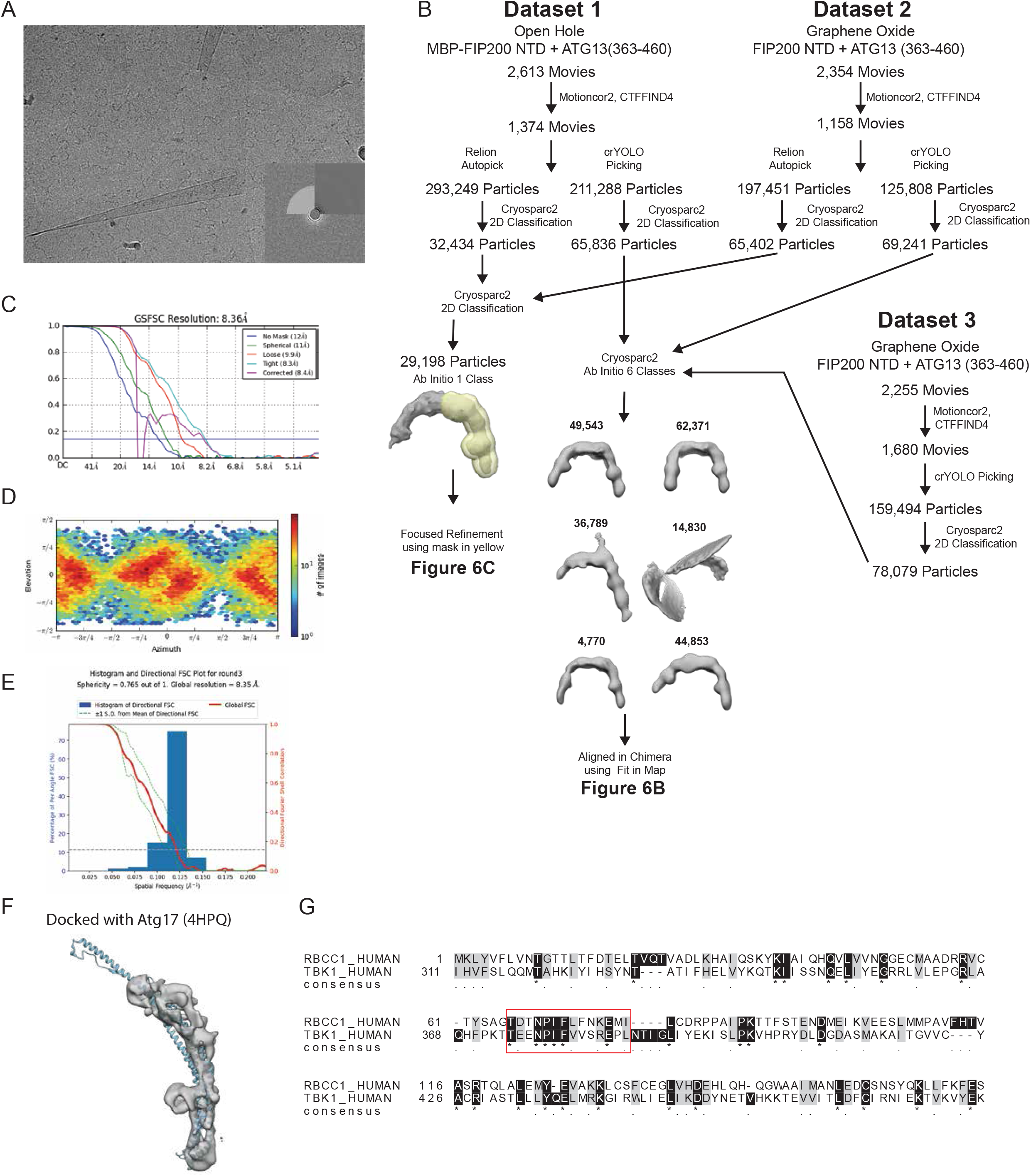
Cryo-EM workflow. (A) Representative micrograph from Dataset 2. (B) Data processing scheme for final cryo EM reconstructions. (C) FSC curves of final cryo EM reconstruction of FIP200 NTD. (D) Orientation parameters of final particle alignments. (E) 3DFSC plot of the final map (F) Fit of Atg17 structure (PDB:4HPQ, blue) into our cryo EM density. (G) Sequence alignment of FIP200 (RBCC1) and human TBK1.

**Table S1.**

| Dataset | FIP200NTD | MBP-FIP200NTD | FIP200NTD-MBP | FIP200NTD, ATG13, ATG101 | MBP-FIP200NTD, ATG13, ATG101 |
| --- | --- | --- | --- | --- | --- |
| Microscope | TF20 |  |  | T12 |  |
| Pixel Size (A/pixel) | 1.5 |  |  | 2.2 |  |
| Micrographs | 734 | 59 | 74 | 325 | 110 |
| Defocus Range | 3.0-4.0 | 1.0-3.0 | 1.0-2.5 | 0.5-4.5 | 0.5-1.5 |
| Box size (unbinned) | 288x288 | 200x200 | 200x200 | 200x200 | 200x200 |
| Particle Picker |  |  | Relion LoG Picker |  |  |
| Particle Picks | 189906 | 42394 | 15446 | 68165 | 44995 |
| Software |  |  | Cryosparc2 |  |  |
| Final Class Avg. # | 80 | 80 | 50 | 50 | 50 |
| Particles in 2D Avgs. | 58001 | 20737 | 7764 | 18417 | 4767 |
| Box Size (2D avgs.) | 144x144 | 100x100 | 100x100 | 100x100 | 100x100 |
| Figure | 1C | 1C | 1C | 2B | 2B |

| Datset | FIP200NTD-MBP, ATG13, ATG101 | FIP200NTD, ATG13(363-460) | FIP200NTD, MBP-ATG13 (363-460) | FIP200 NTD, MBP-ATG13, MBP-ATG101 | FIP200NTD, ATG13-MBP, MBP-ULK1 EAT |
| --- | --- | --- | --- | --- | --- |
| Microscope |  |  | T12 |  |  |
| Pixel Size (A/pixel) |  |  | 2.2 |  |  |
| Micrographs | 127 | 53 | 360 | 62 | 110 |
| Defocus Range | 0.5-1.5 | 0.5-4.5 | 0.5-4.5 | 0.5-2.5 | 0.5-2.5 |
| Box size (unbinned) | 200x200 | 200x200 | 200x200 | 320x320 | 200x200 |
| Particle Picker |  |  | Relion LoG Picker |  |  |
| Particle Picks | 46226 | 18642 | 106018 | 27943 | 21845 |
| Software |  |  | Cryosparc2 |  |  |
| Final Class Avg. # | 50 | 50 | 50 | 50 | 50 |
| Particles in 2D Avgs. | 10954 | 5529 | 24103 | 5090 | 5354 |
| Box Size (2D avgs.) | 100x100 | 100x100 | 100x100 | 160x160 | 100x100 |
| Figure | 2B | 3C | 5A | 5B | 5C |

**Table S3.**
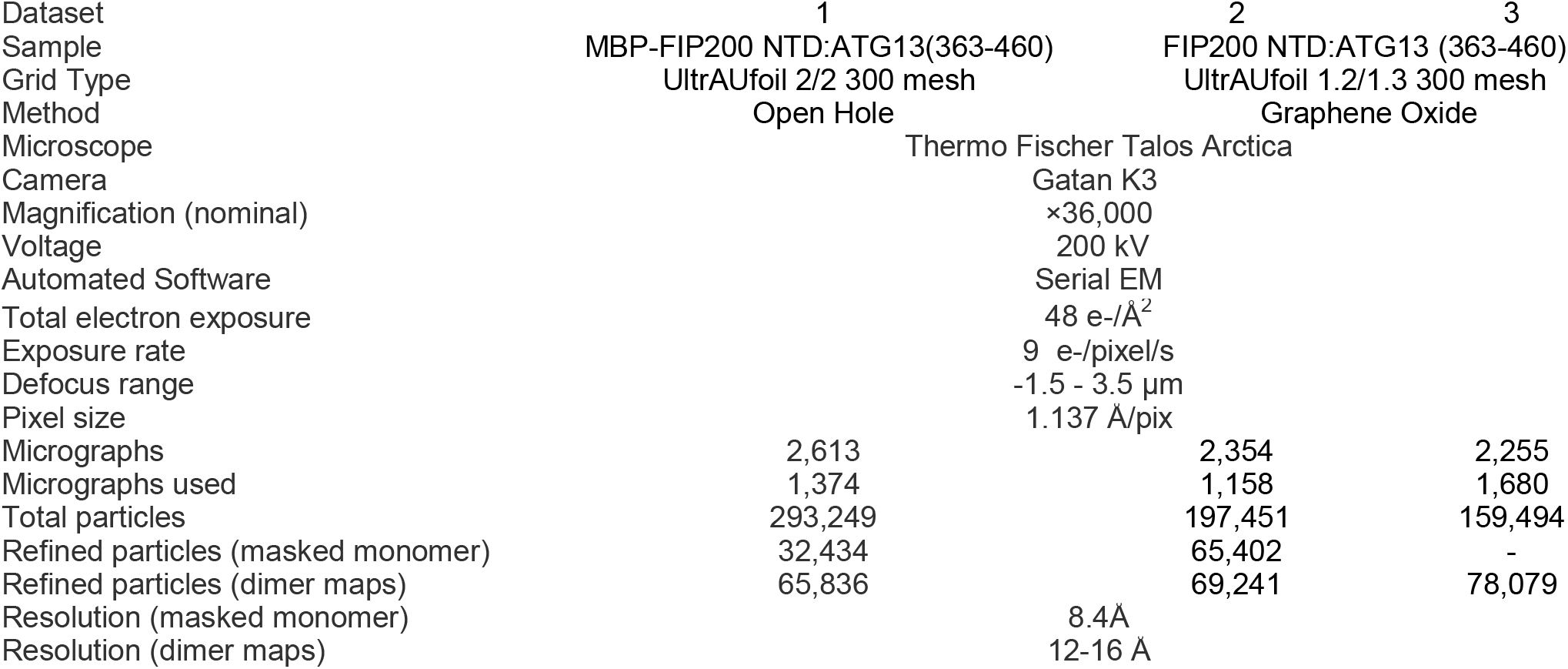
Cryo EM Data Collection and Processing.

**Table S4.**

| Construct | Figure | Source |
| --- | --- | --- |
| GST-TEVcs-FIP200(N-640)-MBP | 1B | This study |
| GST-TEVcs-FIP200(636-C)-MBP | 1B | This study |
| MBP-TSF-TEVcs-ULK1 | 1B, 2A, 4A-C | This study |
| TSF-ATG13 | 1B, 2A-B, 2D-E, S4 | This study |
| TSF-ATG101 | 1B, 2A-C, 2E, 3A-B, 4A-C, S4 | This study |
| GST-TEVcs-FIP200(N-640) | 2B-E, 3A-C, 4A-C, 5A-E, 6, S4, S5 | This study |
| MBP-FIP200(N-640) | 1C, 2B | This study |
| FIP200(N-640)-MBP | 1C, 2B, 3D | This study |
| GST-TEVcs-FIP200(N-640)-Strep | 1C-D, 2A, 2C | This study |
| ATG13 | 2C, 3A-B, 4A-C | This study |
| TSF-TEVcs-ATG13(363-460) | 2C, 3C, 6, S5 | This study |
| ATG101 | 2D, 4A-C, 5C | This study |
| GST-TEVcs-FIP200(N-640)(73-80Mut)M1 | 2D | This study |
| GST-TEVcs-FIP200(N-640)(158-165Mut)M2 | 2D | This study |
| GST-TEVcs-FIP200(N-640)(319-326Mut)M3 | 2D | This study |
| GST-TEVcs-FIP200(N-640)(350-357Mut)M4 | 2D | This study |
| GST-TEVcs-FIP200(N-640)(435-442Mut)M5 | 2D | This study |
| GST-TEVcs-FIP200(N-640)(443-450Mut)M6 | 2D | This study |
| GST-TEVcs-FIP200(N-640)(464-471Mut)M7 | 2D | This study |
| GST-TEVcs-FIP200(N-640)(482-489Mut)M8 | 2D | This study |
| GST-TEVcs-FIP200(N-640)(537-544Mut)M9 | 2D | This study |
| GST-TEVcs-FIP200NTD 4Amut | 2E | Chen *et al.*, *Genes Dev*, 2016 |
| GST-TEVcs-ATG13 | 3A-B | This study |
| GST-TEVcs-ATG13(363-460) | 3D | This study |
| GST-TEVcs-ATG13(363-460)(371-378Mut)M1 | 3D | This study |
| GST-TEVcs-ATG13(363-460)(390-397Mut)M2 | 3D | This study |
| GST-TEVcs-ATG13(363-460)(446-453Mut)M3 | 3D | This study |
| TSF-ULK1(831-C) | 4A-C, 5B | This study |
| MBP-ATG13(363-460) | 5A, 5D | This study |
| MBP-ATG13 | 5B | This study |
| MBP-ATG101 | 5B | This study |
| GST-TEVcs-ATG13-MBP | 5C | This study |
| TSF-ULK1(831-C)-MBP | 5C | This study |
| TSF-TEVcs-ATG13(363-C) | 5E | This study |
| GST-TEVcs-ULK1(831-C)-MBP | 5E | This study |
| MBP-TEVcs-ULK1-6His | S4 | This study |
| MBP-TEVcs-ULK1(N-830)-6His | S4 | This study |
| TSF-ATG13(N-486) | S4 | This study |

## References

Anding, A.L., and E.H. Baehrecke. 2017. Cleaning House: Selective Autophagy of Organelles. Dev. Cell. 41:10–22.

Bahrami, A.H., M.G. Lin, X. Ren, J.H. Hurley, and G. Hummer. 2017. Scaffolding the cup-shaped double membrane in autophagy. PLoS Comput Biol. 13:e1005817.

Behrends, C., M.E. Sowa, S.P. Gygi, and J.W. Harper. 2010. Network organization of the human autophagy system. Nature. 466:68–76.

Bento, C.F., M. Renna, G. Ghislat, C. Puri, A. Ashkenazi, M. Vicinanza, F.M. Menzies, and D.C. Rubinsztein. 2016. Mammalian Autophagy: How Does It Work? Annu. Rev. Biochem. 85:685–713.

Chan, E.Y., A. Longatti, N.C. McKnight, and S.A. Tooze. 2009. Kinase-inactivated ULK proteins inhibit autophagy via their conserved C-terminal domains using an Atg13-independent mechanism. Mol.Cell Biol. 29:157–171.

Chen, S., C. Wang, S. Yeo, C.C. Liang, T. Okamoto, S. Sun, J. Wen, and J.L. Guan. 2016. Distinct roles of autophagy-dependent and -independent functions of FIP200 revealed by generation and analysis of a mutant knock-in mouse model. Genes Dev. 30:856–869.

Fujioka, Y., S.W. Suzuki, H. Yamamoto, C. Kondo-Kakuta, Y. Kimura, H. Hirano, R. Akada, F. Inagaki, Y. Ohsumi, and N.N. Noda. 2014. Structural basis of starvation-induced assembly of the autophagy initiation complex. Nat. Struct. Mol. Biol. 21:513–521.

Ganley, I.G., D.H. Lam, J. Wang, X. Ding, S. Chen, and X. Jiang. 2009. ULK1-ATG13-FIP200 complex mediates mTOR signaling and is essential for autophagy. J. Biol. Chem. 284:12297–12305.

Gomes, L.C., and I. Dikic. 2014. Autophagy in Antimicrobial Immunity. Mol. Cell. 54:224–233.

Hara, T., A. Takamura, C. Kishi, S.I. Iemura, T. Natsume, J.L. Guan, and N. Mizushima. 2008. FIP200, a ULK-interacting protein, is required for autophagosome formation in mammalian cells. J. Cell Biol. 181:497–510.

Hayashi-Nishino, M., N. Fujita, T. Noda, A. Yamaguchi, T. Yoshimori, and A. Yamamoto. 2009. A subdomain of the endoplasmic reticulum forms a cradle for autophagosome formation. Nat.Cell Biol. 11:1433–1437.

Hieke, N., A.S. Loffler, T. Kaizuka, N. Berleth, P. Bohler, S. Driessen, F. Stuhldreier, O. Friesen, K. Assani, K. Schmitz, C. Peter, B. Diedrich, J. Dengjel, P. Holland, A. Simonsen, S. Wesselborg, N. Mizushima, and B. Stork. 2015. Expression of a ULK1/2 binding-deficient ATG13 variant can partially restore autophagic activity in ATG13-deficient cells. Autophagy. 11:1471–1483.

Hosokawa, N., T. Hara, T. Kaizuka, C. Kishi, A. Takamura, Y. Miura, S.I. Iemura, T. Natsume, K. Takehana, N. Yamada, J.L. Guan, N. Oshiro, and N. Mizushima. 2009. Nutrient-dependent mTORC1 Association with the ULK1-Atg13-FIP200 Complex Required for Autophagy. Molecular Biology of the Cell. 20:1981–1991.

Hurley, J.H., and L.N. Young. 2017. Mechanisms of Autophagy Initiation. Annu. Rev. Biochem. 86:225–244.

Itakura, E., and N. Mizushima. 2010. Characterization of autophagosome formation site by a hierarchical analysis of mammalian Atg proteins. Autophagy. 6:764–776.

Jung, C.H., C.B. Jun, S.H. Ro, Y.M. Kim, N.M. Otto, J. Cao, M. Kundu, and D.H. Kim. 2009. ULK-Atg13-FIP200 Complexes Mediate mTOR Signaling to the Autophagy Machinery. Molecular Biology of the Cell. 20:1992–2003.

Kamada, Y., T. Funakoshi, T. Shintani, K. Nagano, M. Ohsumi, and Y. Ohsumi. 2000. Tor-mediated induction of autophagy via an Apg1 protein kinase complex. J. Cell Biol. 150:1507–1513.

Kamada, Y., K. Yoshino, C. Kondo, T. Kawamata, N. Oshiro, K. Yonezawa, and Y. Ohsumi. 2010. Tor Directly Controls the Atg1 Kinase Complex To Regulate Autophagy. Mol. Cell. Biol. 30:1049–1058.

Karanasios, E., E. Stapleton, M. Manifava, T. Kaizuka, N. Mizushima, S.A. Walker, and N.T. Ktsitakis. 2013. Dynamic association of the ULK1 complex with omegasomes during autophagy induction. J. Cell Sci. epub ahead of print.

Karanasios, E., S.A. Walker, H. Okkenhaug, M. Manifava, E. Hummel, H. Zimmermann, Q. Ahmed, M.C. Domart, L. Collinson, and N.T. Ktistakis. 2016. Autophagy initiation by ULK complex assembly on ER tubulovesicular regions marked by ATG9 vesicles. Nature Communications. 7:12420.

Kenny, S.J., Chen, X., Ge, L. and Xu, K. 2019. Super-resolution microscopy unveils FIP200-scaffolded, cup-shaped organization of mammalian autophagic initiation machinery. bioRxiv. dx.doi.org/10.1101/712828.

Koyama-Honda, I., E. Itakura, T.K. Fujiwara, and N. Mizushima. 2013. Temporal analysis of recruitment of mammalian ATG proteins to the autophagosome formation site. Autophagy. 9:epub ahead of print.

Lazarus, M.B., C.J. Novotny, and K.M. Shokat. 2015. Structure of the Human Autophagy Initiating Kinase ULK1 in Complex with Potent Inhibitors. ACS chemical biology. 10:257–261.

Lin, M.G., and J.H. Hurley. 2016. Structure and function of the ULK1 complex in autophagy. Curr. Opin. Cell Biol. 39:61–68.

Mercer, C.A., A. Kaliappan, and P.B. Dennis. 2009. A novel, human Atg13 binding protein, Atg101, interacts with ULK1 and is essential for macroautophagy. Autophagy. 5.

Mercer, T.J., A. Gubas, and S.A. Tooze. 2018. A molecular perspective of mammalian autophagosome biogenesis. J. Biol. Chem. jbc.R117.810366.

Mizushima, N. 2010. The role of the Atg1/ULK1 complex in autophagy regulation. Curr. Opin. Cell Biol. 22:132–139.

Mizushima, N., T. Yoshimori, and Y. Ohsumi. 2011. The role of Atg proteins in autophagosome formation. Annu Rev Cell Dev Biol. 27:107–132.

Papinski, D., and C. Kraft. 2016. Regulation of autophagy by signalling through the Atg1/ULK1 complex. J. Mol. Biol. 428:1725–1741.

Punjani, A., J.L. Rubinstein, D.J. Fleet, and M.A. Brubaker. 2017. cryoSPARC: algorithms for rapid unsupervised cryo-EM structure determination. Nat. Methods. 14:290–296.

Qi, S.Q., D.J. Kim, G. Stjepanovic, and J.H. Hurley. 2015. Structure of the Human Atg13-Atg101 HORMA Heterodimer: an Interaction Hub within the ULK1 Complex. Structure. 23:1848–1857.

Ragusa, M.J., R.E. Stanley, and J.H. Hurley. 2012. Architecture of the Atg17 Complex as a Scaffold for Autophagosome Biogenesis. Cell. 151:1501–1512.

Ravenhill, B.J., K.B. Boyle, N. von Muhlinen, C.J. Ellison, G.R. Masson, E.G. Otten, A. Foeglein, R. Williams, and F. Randow. 2019. The Cargo Receptor NDP52 Initiates Selective Autophagy by Recruiting the ULK Complex to Cytosol-Invading Bacteria. Mol Cell. 74:320–329 e326.

Rohou, A., and N. Grigorieff. 2015. CTFFIND4: Fast and accurate defocus estimation from electron micrographs. J Struct Biol. 192:216–221.

Shu, C., B. Sankaran, C.T. Chaton, A.B. Herr, A. Mishra, J. Peng, and P. Li. 2013. Structural insights into the functions of TBK1 in innate antimicrobial immunity. Structure. 21:1137–1148.

Song, Y., F. DiMaio, R.Y. Wang, D. Kim, C. Miles, T. Brunette, J. Thompson, and D. Baker. 2013. High-resolution comparative modeling with RosettaCM. Structure. 21:1735–1742.

Stjepanovic, G., C.W. Davies, R.E. Stanley, M.J. Ragusa, D.J. Kim, and J.H. Hurley. 2014. Assembly and dynamics of the autophagy initiating Atg1 complex. Proc Natl Acad Sci U S A. 111:12793–12798.

Suzuki, H., T. Kaizuka, N. Mizushima, and N.N. Noda. 2015. Structure of the Atg101-Atg13 complex reveals essential roles of Atg101 in autophagy initiation. Nat. Struct. Mol. Biol. 22:572–580.

Turco, E., M. Witt, C. Abert, T. Bock-Bierbaum, M.Y. Su, R. Trapannone, M. Sztacho, A. Danieli, X. Shi, G. Zaffagnini, A. Gamper, M. Schuschnig, D. Fracchiolla, D. Bernklau, J. Romanov, M. Hartl, J.H. Hurley, O. Daumke, and S. Martens. 2019. FIP200 Claw Domain Binding to p62 Promotes Autophagosome Formation at Ubiquitin Condensates. Mol Cell. 74:330–346 e311.

Vargas, J.N.S., C. Wang, E. Bunker, L. Hao, D. Maric, G. Schiavo, F. Randow, and R.J. Youle. 2019. Spatiotemporal Control of ULK1 Activation by NDP52 and TBK1 during Selective Autophagy. Mol Cell. 74:347–362 e346.

Wagner, T., F. Merino, M. Stabrin, T. Moriya, C. Antoni, A. Apelbaum, P. Hagel, O. Sitsel, T. Raisch, D. Prumbaum, D. Quentin, D. Roderer, S. Tacke, B. Siebolds, E. Schubert, T.R. Shaikh, P. Lill, C. Gatsogiannis, and S. Raunser. 2019. SPHIRE-crYOLO is a fast and accurate fully automated particle picker for cryo-EM. Communications biology. 2:218.

Wallot-Hieke, N., N. Verma, D. Schlutermann, N. Berleth, J. Deitersen, P. Bohler, F. Stuhldreier, W. Wu, S. Seggewiss, C. Peter, H. Gohlke, N. Mizushima, and B. Stork. 2018. Systematic analysis of ATG13 domain requirements for autophagy induction. Autophagy. 14:743–763.

Wen, X., and D.J. Klionsky. 2016. An overview of macroautophagy in yeast. J. Mol. Biol. 428:1681–1699.

Yla-Anttila, P., H. Vihinen, E. Jokita, and E.-L. Eskelinen. 2009. 3D tomography reveals connections between the phagophore and endoplasmic reticulum. Autophagy. 5:1180–1185.

Yorimitsu, T., and D.J. Klionsky. 2005. Atg11 links cargo to the vesicle-forming machinery in the cytoplasm to vacuole targeting pathway. Molecular Biology of the Cell. 16:1593–1605.

Zachari, M., and I.G. Ganley. 2017. The mammalian ULK1 complex and autophagy initiation. Essays in biochemistry. 61:585–596.

Zaffagnini, G., and S. Martens. 2016. Mechanisms of selective autophagy. J. Mol. Biol. 428:1714–1724.

Zhang, C., G. Shang, X. Gui, X. Zhang, X.C. Bai, and Z.J. Chen. 2019. Structural basis of STING binding with and phosphorylation by TBK1. Nature. 567:394–398.

Zheng, S.Q., E. Palovcak, J.P. Armache, K.A. Verba, Y. Cheng, and D.A. Agard. 2017. MotionCor2: anisotropic correction of beam-induced motion for improved cryo-electron microscopy. Nat Methods. 14:331–332.

Zivanov, J., T. Nakane, B.O. Forsberg, D. Kimanius, W.J. Hagen, E. Lindahl, and S.H. Scheres. 2018. New tools for automated high-resolution cryo-EM structure determination in RELION-3. Elife. 7.

